# HIV-1 vaccine design through minimizing envelope metastability

**DOI:** 10.1101/361931

**Authors:** Linling He, Sonu Kumar, Joel D. Allen, Deli Huang, Xiaohe Lin, Colin J. Mann, Karen L. Saye-Francisco, Jeffrey Copps, Anita Sarkar, Gabrielle S. Blizard, Gabriel Ozorowski, Devin Sok, Max Crispin, Andrew B. Ward, David Nemazee, Dennis R. Burton, Ian A. Wilson, Jiang Zhu

## Abstract

Overcoming envelope metastability is crucial to trimer-based HIV-1 vaccine design. Here, we present a coherent vaccine strategy by minimizing metastability. For ten strains across five clades, we demonstrate that gp41 ectodomain (gp41_ECTO_) is the main source of envelope metastability by replacing wild-type gp41_ECTO_ with BG505 gp41_ECTO_ of the uncleaved prefusion-optimized (UFO) design. These gp41_ECTO_-swapped trimers can be produced in CHO cells with high yield and high purity. Crystal structure of a gp41_ECTO_-swapped trimer elucidates how a neutralization-resistant tier 3 virus evades antibody recognition of the V2 apex. UFO trimers of transmitted/founder (T/F) viruses and UFO trimers containing a consensus-based ancestral gp41_ECTO_ suggest an evolutionary root of the metastability. Gp41ECTO-stabilized trimers can be readily displayed on 24- and 60-meric nanoparticles, with incorporation of additional T cell help illustrated for a hyperstable 60-mer. In mice and rabbits, gp140 nanoparticles induced more effective tier 2 neutralizing antibody response than trimers with statistical significance.

**HIGHLIGHTS:**

- gp41 is the main source of HIV-1 envelope metastability
- BG505 gp41 of the UFO design stabilizes gp140 trimers of diverse subtypes
- gp41 stabilization facilitates gp140 nanoparticle assembly and improves production
- Nanoparticles elicit tier 2 neutralizing antibodies more effectively than trimers

## INTRODUCTION

The envelope glycoprotein (Env) of human immunodeficiency virus type-1 (HIV-1) harbors the epitopes of all broadly neutralizing antibodies (bNAbs) (Burton and Mascola, 2015) and is the main target of vaccine design (Haynes and Mascola, 2017). The cleaved, mature Env is presented on HIV-1 virions as a metastable trimer of heterodimers each containing a (co-) receptor-binding protein, gp120, and a transmembrane protein, gp41, which anchors the Env spike within the viral membrane and drives the fusion process during cell entry (Wyatt and Sodroski, 1998). Due to its labile nature and a dense layer of surface glycans (Scanlan et al., 2007), Env has long resisted structure determination and trimer-based vaccine design efforts. While the functional necessity of a metastable Env for HIV-1 infection is well understood, the molecular source of metastability and how to eliminate it from the Env trimer remain unclear. It is perhaps not an overstatement that overcoming Env metastability has been and still is central to trimer-based HIV-1 vaccine design.

Several strategies have been proposed to create stable and soluble gp140 trimers as potential vaccine immunogens. The early generation of gp140 trimers was based on intuitive designs to overcome metastability via deletion of the cleavage site between gp120 and the gp41 ectodomain (gp41_ECTO_) and the addition of trimerization motifs at the C terminus (Yang et al., 2000a; Yang et al., 2000b; Yang et al., 2002). A more rigorous trimer design, designated SOSIP.664, stresses the importance of cleavage, trimer stability and solubility (Sanders et al., 2002). This format utilizes a disulfide bond between gp120 and gp41_ECTO_, an I559P mutation in the heptad repeat 1 (HR1) region, and a truncation at position 664. When applied to clade A BG505, the SOSIP trimer was shown to be a close mimic of the native Env spike (Sanders et al., 2013) that enabled Env structure determination for the first time by X-ray crystallography and electron microscopy (EM) (Julien et al., 2013a; Lyumkis et al., 2013). The SOSIP trimer has significantly advanced HIV-1 research by providing an antigen for bNAb isolation and structural characterization (Blattner et al., 2014; Doria-Rose et al., 2016; Falkowska et al., 2014; Garces et al., 2015; Garces et al., 2014; Huang et al., 2014; Kong et al., 2016b; Lee et al., 2017; Lee et al., 2016; Ozorowski et al., 2017; Scharf et al., 2014; Sok et al., 2014), a strategy for stabilizing diverse Envs (de Taeye et al., 2015; Julien et al., 2015; Pugach et al., 2015; Stewart-Jones et al., 2016), and a template for trimer optimization either to shield non-neutralizing antibody (non-NAb) epitopes or to target bNAb precursors (Chuang et al., 2017; Do Kwon et al., 2015; Ringe et al., 2017a; Steichen et al., 2016). As the SOSIP design continued to evolve, new versions (designated SOSIP.v5 and v6) were proposed that further improved trimer stability and immunogenicity (de la Pena et al., 2017). Removal of the cleavage site has proven successful in the forms of single-chain gp140 (sc-gp140) (Georgiev et al., 2015), native flexibly linked (NFL) (Guenaga et al., 2015; Sharma et al., 2015), and uncleaved prefusion-optimized (UFO) (Kong et al., 2016a) trimers. However, it was not until recently that the primary cause of Env metastability – an HR1 bend (residues 547-569) in gp41_ECTO_ – was identified and targeted directly by rational redesign (Kong et al., 2016a). Of note, the I559P mutation that improved trimer stability in the SOSIP design is within this HR1 bend (Sanders et al., 2013). Glycine substitution in this region was also found to improve trimer properties in the NFL form (Guenaga et al., 2017). While attempts to elicit tier 2 NAb response with BG505 SOSIP trimers in wild-type (WT) mice were unsuccessful (Hu et al., 2015), both SOSIP and NFL trimers induced consistent autologous tier 2 NAbs in rabbits and non-human primates (NHPs), with sporadic neutralization observed for heterologous tier 2 isolates (de Taeye et al., 2015; Klasse et al., 2016; Martinez-Murillo et al., 2017; McCoy et al., 2016; Pauthner et al., 2017; Sanders et al., 2015). Therefore, despite recent advances in rational design and structural analysis of native-like trimers (Sanders and Moore, 2017; Ward and Wilson, 2017), significant barriers still remain in the path to an effective HIV-1 vaccine.

In this study, we set out to address the diverse challenges in HIV-1 vaccine development with a coherent strategy centered on Env metastability. We first examined the utility of a transient CHO cell line (ExpiCHO) to express native-like trimers, which resulted in superior trimer yield, purity, and antigenicity while displaying subtle differences in glycosylation pattern and B cell response compared to 293 F-produced trimers. We then demonstrated that gp41_ECTO_ is the main source of metastability by replacing WT gp41_ECTO_ with BG505 gp41_ECTO_ of the UFO design for ten Envs across clades A, B, C, B/C, and A/E. The gp41_ECTO_-swapped trimers (termed UFO-BG) exhibited substantial yield and purity and were structurally validated by negative-stain EM. Analysis of UFO and UFO-BG trimers by bio-layer interferometry (BLI) against a large panel of nineteen antibodies provided comprehensive antigenic profiles for ten Envs across five subtypes. Crystal structure was determined for the H078.14 UFO-BG trimer (tier 3, clade B) at 4.4 Å resolution, which explains how this virus evades bNAbs that target the apex and enables antigenic optimization of this tier 3 trimer. We also observed high yield, high purity, and native-like antigenicity for the UFO trimers of transmitted/founder (T/F) viruses and the UFO trimers containing a consensus gp41_ECTO_ (termed UFO-C), suggesting an evolutionary explanation of Env metastability. Next, we displayed diverse UFO-BG trimers on a 24-meric ferritin nanoparticle (He et al., 2016) and demonstrated how to incorporate T cell help into nanoparticle constructs for a hyperstable 60-mer, I3-01 (Hsia et al., 2016). In WT mice, ferritin and I3-01 nanoparticles, as well as a scaffolded gp140 trimer, induced autologous tier 2 NAbs to BG505.T332N after eight weeks, whereas soluble trimers did not. In rabbits, the ferritin nanoparticle elicited an autologous tier 2 NAb response after six weeks, which was statistically different than the tier 2 NAb response elicited by the soluble trimer throughout the course of immunization. Our study thus presents a coherent strategy for HIV-1 vaccine design along with vaccine candidates that merit further evaluation in NHPs and potentially in humans.

## RESULTS

### A Robust Expression System for Production of Native-Like gp140 Trimers

Rapid growth in protein therapeutics and vaccines has accelerated the development of high-yield mammalian cell lines (Chu and Robinson, 2001). In the past, native-like trimers were primarily characterized in laboratory expression systems, with uncertainties in their transferability to an industrial setting. The CHO cell line is one of the principal mammalian expression systems that meet the Good Manufacturing Practice (GMP) standard. Although SOSIP and NFL trimers have been produced in stable CHO cell lines for *in vitro* and *in vivo* testing, gp140 modification and bNAb affinity purification were required to achieve high-quality materials (Chung et al., 2014; Dey et al., 2017; Pauthner et al., 2017; Ringe et al., 2017b). A transient, high-yield CHO cell line derived from the GMP CHO-S cells, such as ExpiCHO, would likely accelerate the evaluation of new trimer designs and their development towards vaccine candidates.

Previously, we reported ExpiCHO expression of gp140-ferritin nanoparticles (He et al., 2016). Here, we examined the utility of ExpiCHO for producing native-like Env trimers. First, we transiently expressed BG505 SOSIP, HR1-redesigned, and UFO gp140 trimers in ExpiCHO cells (all gp140 constructs tested hereafter were truncated at residue 664 unless stated otherwise). Env proteins were extracted from the supernatants using a *Galanthus nivalis* lectin (GNL) column and purified by size exclusion chromatography (SEC) on a Superdex 200 16/600 column. Ultraviolet absorbance at 280 nm (UV_280_) was utilized as a metric to compare the SEC profiles (Figure 1A). Remarkably, a 100-ml ExpiCHO expression produced well-folded gp140 protein equivalent to that obtained from 2-4 liters of 293 F cells (5-12 mg prior to SEC). Overall, the SEC profiles exhibited a significant reduction of misfolded species in the Env protein produced by ExpiCHO cells as compared to 293 F cells (Kong et al., 2016a). While the SOSIP and HR1-redesigned trimers were still mixed with small amounts of aggregates, as shown by the shoulder to the left of the trimer peak at 55 ml, the UFO trimer displayed a single peak indicative of homogeneity. In addition, the UV_280_ value of the UFO trimer was 2.6 and 1.2-fold greater than that of the SOSIP and HR1-redesigned trimers, respectively, indicating a higher yield for the UFO trimer. Blue native polyacrylamide gel electrophoresis (BN-PAGE) showed a characteristic trimer band across all SEC fractions with minimal impurity (Figure 1B). We then evaluated trimer antigenicity using BLI and a panel of six bNAbs and four non-NAbs (Figure 1C and Figure S1A). While binding kinetics appeared to be largely independent of the cell lines used, trimers produced in ExpiCHO cells showed enhanced bNAb recognition relative to those in 293 F cells (Kong et al., 2016a). Among the three trimer constructs, UFO displayed the least binding to non-NAbs, consistent with its high purity and stability, although all three trimers bound to a V3-specific non-NAb, 19b.

**Figure 1.**
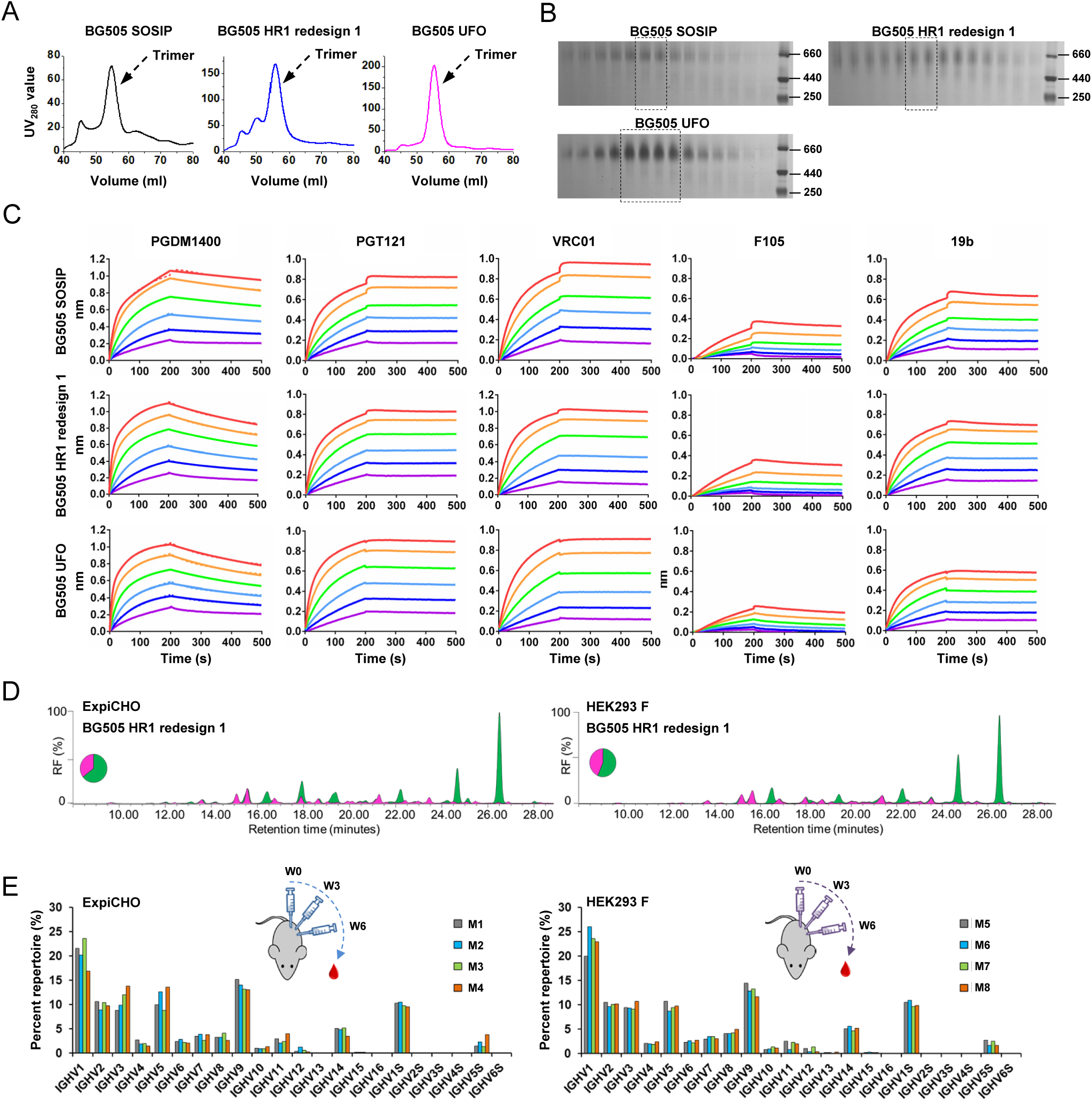
Characterization of ExpiCHO-produced native-like Env trimers. (**A**) SEC profiles of ExpiCHO-expressed, GNL-purified BG505 SOSIP.664 trimer, HR1-redesigned trimer (HR1 redesign 1), and UFO trimer from a Superdex 200 16/600 column. Transient expression in 200 ml ExpiCHO cells was used. (**B**) BN-PAGE of Env proteins for the three abovementioned trimers. The fractions used for antigenic profiling are circled by black dotted lines on the gel. (**C**) Antigenic profiles of the purified trimers measured against a panel of representative bNAbs and non-NAbs, with additional antibody binding profiles shown in Figure Sl. Sensorgrams were obtained on an Octet RED96 using a trimer titration series of six concentrations (200-6.25 nM by two-fold dilution). (**D**) HILIC-UPLC profiles of the enzymatically released A-linked glycans of the HR1-redesigned trimer produced in ExpiCHO and 293 F cells followed by GNL and SEC purification. Oligomannose-type and hybrid glycans (green) were identified by their sensitivity to endo H digestion. Peaks corresponding to complex-type glycans are shown in pink. The peaks are integrated, and the pie charts summarize the quantification of the peak areas. (**E**) Germline gene usage of the mouse antibody repertoire primed by ExpiCHO and 293 F-expressed BG505 gp140 trimers containing the HR1 redesign. The percentage of each germline gene family is plotted as a histogram, with a cartoon picture showing the immunization scheme. The four mice in each group are colored in gray (M1), cyan (M2), light green (M3), and orange (M4).

Next, we investigated the *N*-glycan processing of the cleaved HR1-redesigned BG505 trimer (Kong et al., 2016a) in ExpiCHO and 293 F cells using methods previously reported for the cleaved BG505 SOSIP trimer (Behrens et al., 2017; Behrens et al., 2016). The oligomannose content of gp140 was quantified using ultra-high-performance liquid chromatography (UPLC) (Figure 1D and Figure S1B). Glycans released from 293 F-expressed gp140 consisted of 56% oligomannose-type and 44% complex-type, while a slightly elevated proportion of oligomannose-type glycans, 64%, was observed for the same construct expressed in ExpiCHO cells. The oligomannose content of gp140 expressed in both cell lines appeared to be similar to that observed for the BG505 SOSIP trimer, approximately 63% (Behrens et al., 2016) or 68% (Dey et al., 2017). Site-specific glycan analysis was performed using liquid chromatography–mass spectrometry (LC-MS) to determine the relative intensities of the glycans at each *N*-glycosylation site, revealing features characteristic of the native-like trimers (Figure S1C). As determined by UPLC, many glycosylation sites which present solely oligomannose-type glycans are from regions of underprocessed glycans previously characterized for the SOSIP trimer. For example, glycans at sites N332 and N295 are exclusively oligomannose-type (Behrens et al., 2016), and are the key elements of a glycan supersite targeted by multiple classes of bNAbs recognizing Man9GlcNAc_2_ (Walker et al., 2011). Our analysis thus confirmed that this glycan supersite is present on the HR1-redesigned gp140 trimers produced in both cell lines. Similarly, another oligomannose-rich region encompassing N156 and N160 near the trimer apex showed patterns consistent with those observed for the SOSIP trimer (Behrens et al., 2017; Behrens et al., 2016). Overall, the HR1-redesigned gp140 trimers expressed in either 293 F or ExpiCHO cells presented glycosylation patterns consistent with correctly folded Env. However, some relatively subtle ExpiCHO-specific glycan patterns were also observed, such as a lower to non-detectable proportion of complex-type glycans at N339, which may contribute to the enhanced bNAb binding to the N332 supersite (Figure S1D). To examine whether the cell line-specific glycan patterns affect trimer-induced B cell responses *in vivo*, we immunized BALB/c mice with intraperitoneal (i.p.) injections of 50 μg trimer protein (HR1-redesigned BG505 gp140 trimer) adjuvanted with AddaVax at weeks 0, 3 and 6, and then probed the peripheral B cell repertoires by next-generation sequencing (NGS) (Morris et al., 2017). Although the two groups of mice displayed similar patterns of germline gene usage, slightly increased IGHV3 and IGHV5 frequencies (up to 4%) were observed in the B cell repertoires primed by the ExpiCHO-expressed trimer, suggesting a potential glycan influence on trimer-induced B cell response (Figure 1E).

Taken together, ExpiCHO provides a robust expression system for producing both native-like trimers and gp140 nanoparticles, with important implications for manufacture. In addition to the proper proteolytic cleavage by furin co-expressed in ExpiCHO cells, as previously reported for a standard CHO cell line (Chung et al., 2014), subtle differences in glycan processing and antibody repertoire response are noted between trimers produced in ExpiCHO and 293 F cells.

### Design and Characterization of UFO-BG Trimers for Five HIV-1 Clades

A major obstacle faced by current trimer designs is measurable loss in yield, purity, and stability once they are extended from BG505 to other strains. The solutions proposed thus far include: (1) purification methods aimed to separate native-like trimers from misfolded and other Env species (gp140 monomers, dimers, and aggregates), such as bNAb affinity columns (Pugach et al., 2015), negative selection (Sharma et al., 2015), multi-cycle SEC (Georgiev et al., 2015), and a combined chromatographic approach (Verkerke et al., 2016); and (2) auxiliary Env-stabilizing mutations informed by atomic structures (Chuang et al., 2017; Guenaga et al., 2017) or selected from large-scale screening (Steichen et al., 2016; Sullivan et al., 2017). However, these solutions are empirical by nature and can result in suboptimal outcomes such as reduced trimer yield and bNAb recognition. Recently, we identified an HR1 bend (residues 547-569) in gp41_ECTO_ as the primary cause of Env metastability (Kong et al., 2016a). Rational redesign of this HR1 bend notably improved trimer yield and purity for multiple HIV-1 strains, yet still produced varying amounts of misfolded Env (Kong et al., 2016a). These data suggested that other regions besides HR1, which might be in gp120 and/or gp41_ECTO_, also contribute to the Env metastability. Thus, determining the location of these ‘secondary factors of metastability’ and eliminating them from the Env trimer may prove crucial for trimer-based vaccine design.

Here, we hypothesized that all factors of Env metastability are encoded within gp41_ECTO_, and that BG505 gp41_ECTO_ of the UFO design (termed UFO-BG) can be used to stabilize diverse HIV-1 Envs (Figure 2A). To investigate this hypothesis, we selected ten Envs across five clades (A, B, C, B/C, and A/E) from either a large panel of tiered HIV-1 pseudoviruses (Seaman et al., 2010) or the available database (https://www.hiv.lanl.gov), and included three Envs tested in our previous study (Kong et al., 2016a). Of note, seven of the ten Envs here were derived from tier 2/3 isolates. For each Env, the gp140 constructs of SOSIP, UFO, and UFO-BG designs were transiently expressed in 100-ml ExpiCHO cells, with the SOSIP trimer co-transfected with furin. Following GNL purification, the SEC profiles of thirty gp140s were generated from a Superdex 200 16/600 column for comparison (Figure 2B). In terms of trimer yield, UFO-BG was up to 5- and 53-fold more than UFO and SOSIP, respectively. For all ten Envs, except for BG505, SOSIPs showed a significant proportion of aggregates (at volumes 40-50 ml in the SEC profile) accompanied by an extremely low yield and sometimes absence of a trimer peak. UFOs improved trimer yield and purity considerably, most notably for two clade C strains, though not for clade A/E. The UFO-BG design demonstrated unparalleled trimer yield and purity for eight of ten strains, with no or only slight hints of dimers and monomers. All thirty gp140 proteins were then characterized by BN-PAGE (Figure S2A). Overall, UFO-BG dramatically reduced the dimer and monomer content with respect to SOSIP and UFO, showing a trimer band across all SEC fractions and only occasionally faint bands of lower molecular weight. Based on this finding, we compared the total Env protein obtained from a GNL column against the pooled trimer protein after SEC and fraction analysis. Surprisingly, GNL purification alone yielded comparable purity for all UFO-BG trimers, except for those derived from a tier 2 clade B strain and a tier 3 clade B/C strain (Figure 2C). Next, thermal stability was assessed for eight purified UFO-BG trimers using differential scanning calorimetry (DSC) (Figure 2D and Figure S2B). Notably, the DSC profiles exhibited a clade or strain-specific pattern, with the thermal denaturation midpoint (T_m_) ranging from 60.9°C to 68.4 °C. Among the eight UFO-BG trimers tested, BG505 yielded the highest T_m_ (68.4 °C), which was followed by two clade C trimers (65.2-66.2°C). In the absence of additional disulfide bonds and cavity-filling mutations, the DSC data largely reflected the thermal stability of native Env. It should be noted that the CN54 UFO and UFO-BG constructs contained fourteen mutations (CN54M14), which reduced aggregates for 293 F-produced trimers (Figure S2C). Four UFO-BG trimers were randomly selected from clades B, C, and B/C for expression in 293 F cells and SEC characterization (Figure S2D). UFO-BG was found to improve trimer properties regardless of the expression system but achieved optimal purity when produced in ExpiCHO cells.

**Figure 2.**
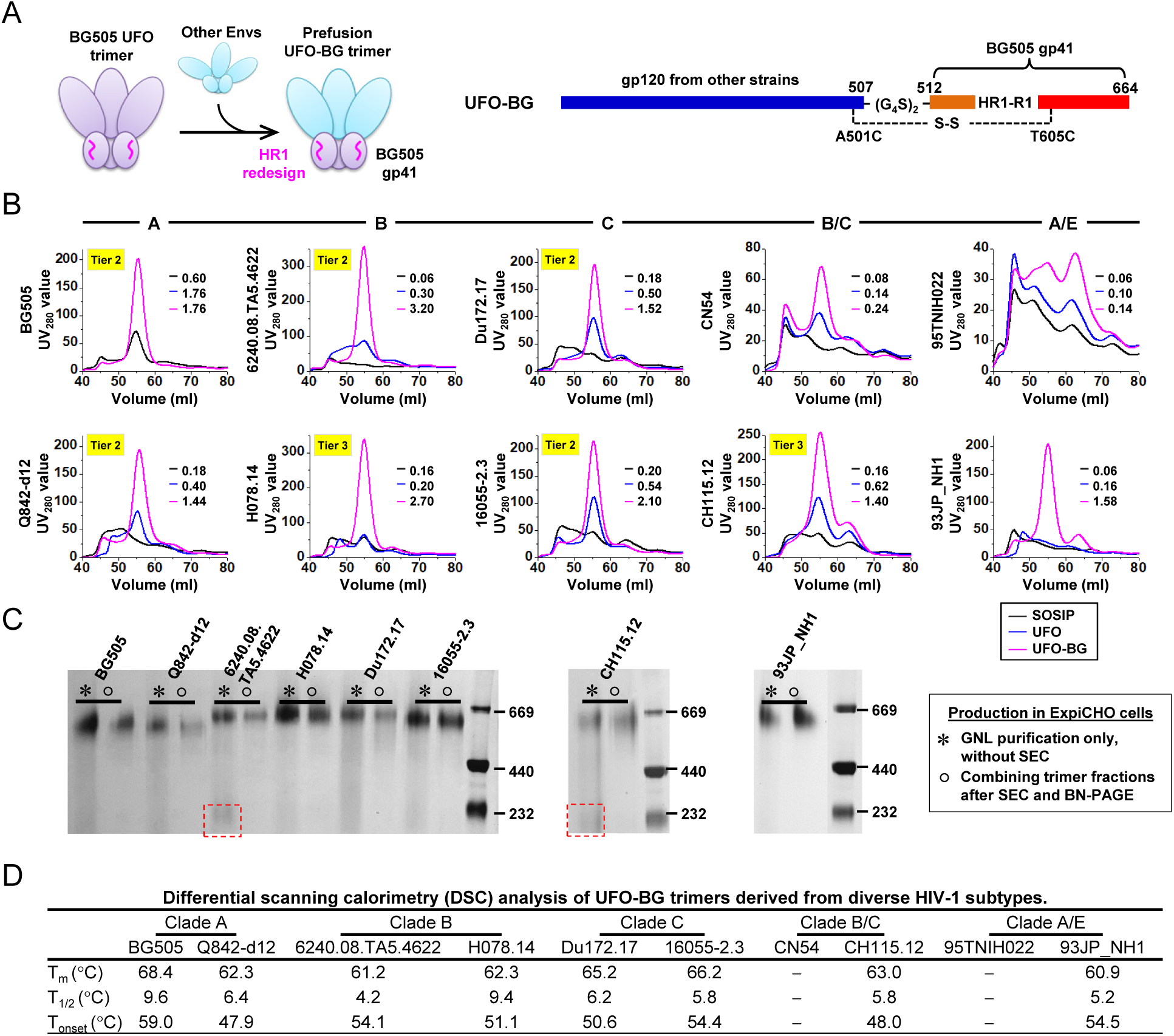
Biochemical and biophysical characterization of the UFO-BG trimers for diverse HIV-1 strains. (**A**) Design (left) and schematic representation (right) of the UFO-BG trimers. As shown on the left, BG505 gp41_ECTO_ of the UFO design is used to stabilize gp120s from other HIV-1 Envs in a hybrid form of gp140 trimer designated UFO-BG. The redesigned HR1 bend is highlighted in magenta. (**B**) SEC profiles of SOSIP, UFO, and UFO-BG trimers derived from ten Envs across five subtypes (A, B, C, B/C, and A/E) following 100-ml ExpiCHO expression and GNL purification. The yield (mg) of SEC-purified trimer protein (fractions corresponding to 5357ml) obtained from 100-ml ExpiCHO expression is listed for each of the three trimer designs (SOSIP, UFO, and UFO-BG). (**C**) BN-PAGE of Env proteins after GNL purification but prior to SEC and of purified trimers following SEC and BN-PAGE for eight UFO-BG constructs. Two recombinant strains, B/C CN54 and A/E 95TNIH022, were not included due to low purity. (**D**) DSC analysis of eight UFO-BG trimers following GNL and SEC purification. Three thermal parameters (Tm, T_1/2_, and T_onset_) are listed for each trimer construct.

The results thus confirm that gp41_ECTO_ is the primary source of metastability and replacement by BG505 gp41_ECTO_ of the UFO design can stabilize diverse Envs. Of note, Env stabilization by BG505 gp41_ECTO_ of the SOSIP design was recently reported (Ahmed et al., 2017; Joyce et al., 2017), but the trigger of Env metastability – the HR1 bend (Kong et al., 2016a) – is still present in the resulting trimers. For UFO-BG trimers, the similar purity before and after SEC suggests the potential for a simple and cost-effective manufacturing solution. The inherent high purity of UFO-BG trimers should also accelerate the development and clinical testing of nucleic acid vaccine strategies (Liu, 2011; Robert-Guroff, 2007; Ulmer et al., 2012).

### Structural Characterization of the UFO-BG Trimers

While biochemical and biophysical properties are informative, structures would provide the most convincing evidence that the UFO-BG trimers are an accurate mimic of the native Env (Ward and Wilson, 2017). The clade B H078.14 gp140 construct was selected for crystallization screening, as the trimer structure for a tier 3 neutralization-resistant isolate has yet to be determined. Briefly, this UFO-BG trimer was produced in 293 F cells with kifunensine treatment to inhibit the complex-type glycan formation, and then purified with a 2G12 affinity column followed by SEC on a Superdex 200 16/600 column. Co-crystallization with antigen-binding fragments (Fabs) of bNAbs PGT124 and 35O22 resulted in a complex structure at a resolution of 4.4 Å (Figure 3A, left, and Figure S3A). Overall, the H078.14 UFO-BG trimer adopts a native-like Env conformation closely resembling that of BG505 SOSIP (PDB: 5CEZ, 3.03Å) and HR1-redesigned (PDB: 5JS9, 6.92Å) trimers (Garces et al., 2015; Kong et al., 2016a), with global C_α_ RMSDs of 0.36 Å and 1.09 Å, respectively (Figure 3A, middle). Small differences were noted at the HR1 bend and the first turn of HR1 central C-terminal helix after structural superposition of gp41_ECTO_ (Figure 3A, right). X-ray crystallographic analysis at this moderate resolution thus confirmed that BG505 gp41_ECTO_ of the UFO design – with a minimal level of metastability – can be used to stabilize diverse HIV-1 Envs in a prefusion state, in addition to providing the first structural model for a tier-3 neutralization-resistant Env spike. Negative-stain EM was utilized to characterize eight UFO-BG trimers which showed substantial purity in SEC and BN-PAGE (Figures 2B and 2C). As indicated by the 2D class averages, 67-100% of GNL-purified Env appeared to be native-like trimers (Figure 3B and **Figure S3B**). Similar results were reported for the SOSIP trimers only after purification using bNAb affinity columns (de la Pena et al., 2017; de Taeye et al., 2015; Julien et al., 2015; Julien et al., 2013b; Pugach et al., 2015; Sanders et al., 2013). Crystallographic and EM analyses thus validated the structural integrity of UFO-BG trimers derived from five subtypes, supporting the notion that UFO-BG is a general and effective strategy for trimer stabilization.

**Figure 3.**
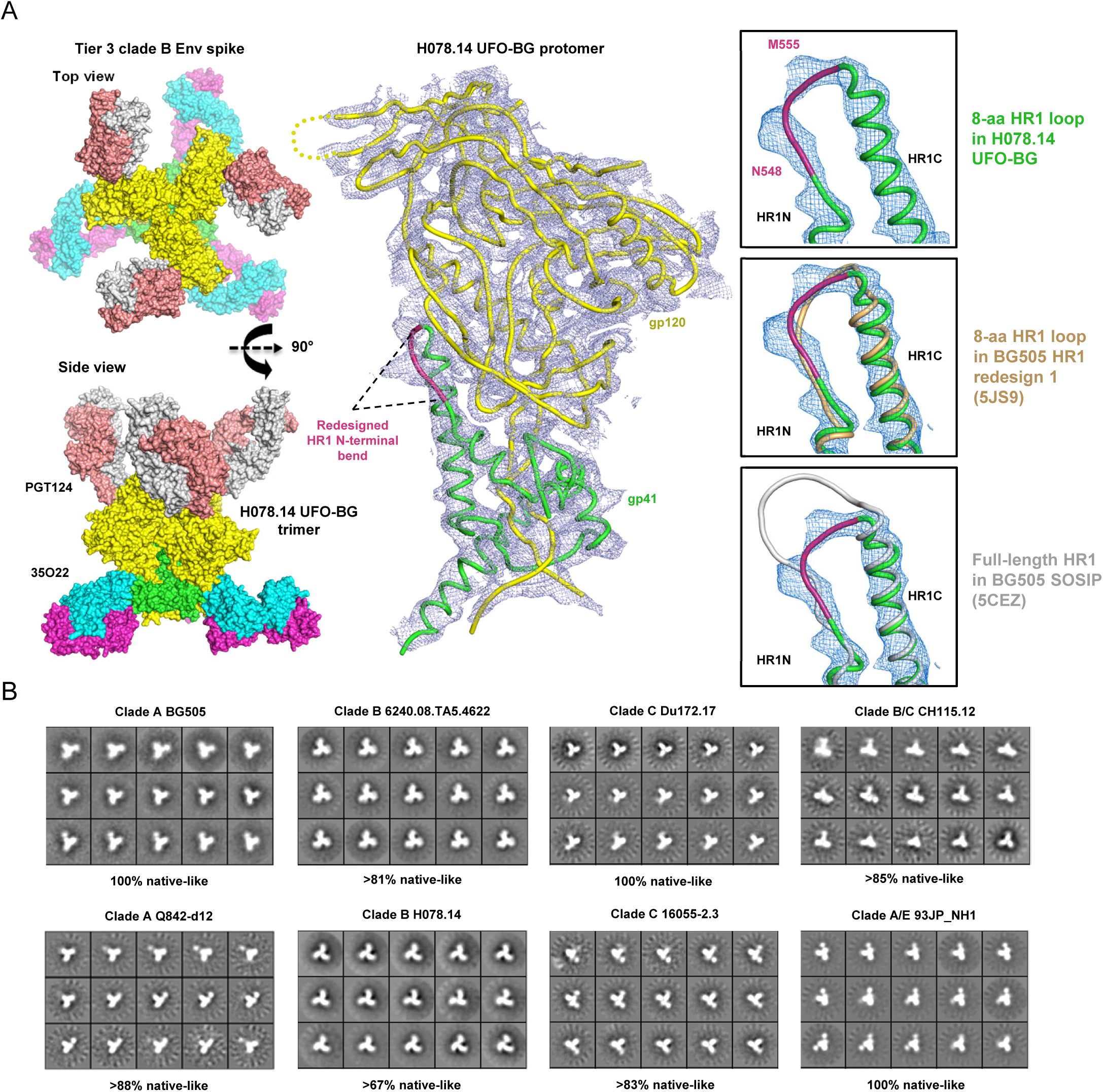
Structural characterization of the UFO-BG trimers derived from diverse HIV-1 Envs. (**A**) Crystal structure of a clade B tier 3 H078.14 Env spike determined at 4.4 Å resolution. Molecular surface of the H078.14 UFO-BG trimer in complex with bNAb Fabs PGT124 and 35O22 is shown on the left (top view and side view), with a ribbon model of the gp140 protomer and two Fabs shown in the middle, and a zoomed-in view of the redesigned HR1 bend (alone and superimposed onto two available structures, 5JS9 and 5CEZ) on the right. (**B**) Reference-free 2D class averages derived from negative-stain EM of eight UFO-BG trimers produced in ExpiCHO cells followed by GNL and SEC purification, with the full sets of images shown in Figure S3B. Percentage of native-like trimers is indicated for each trimer construct.

### Antigenic Evaluation of UFO and UFO-BG Trimers

Following structural characterization, the effect of gp41_ECTO_ substitution on trimer antigenicity was assessed by BLI (Figure 4A and **Figure S4**). To this end, we compared the UFO-BG trimers and UFO trimers containing WT gp41_ECTO_ and a generic GS linker at the HR1 bend (Kong et al., 2016a). Trimer proteins obtained from GNL and SEC purification were tested for antibody binding activity on Octet RED96 as previously described (Kong et al., 2016a). A panel of eleven bNAbs was utilized to assess conserved neutralizing epitopes on the trimer surface, including the V2 apex recognized by PGDM1400 (Sok et al., 2014), PGT145 (Walker et al., 2011), and PG16 (Walker et al., 2009), the N332 supersite by PGT121, PGT128, PGT135 (Walker et al., 2011), and 2G12 (Trkola et al., 1995), the CD4-binding site (CD4bs) by VRC01 (Wu et al., 2010) and b12 (Burton et al., 1994), and the gp120-gp41 interface by PGT151 (Falkowska et al., 2014) and 35O22 (Huang et al., 2014), along with eight non-NAbs targeting the CD4bs, the CD4-induced (CD4i) epitope, the immunodominant epitopes at the V3 tip and within gp41_ECTO_.

**Figure 4.**
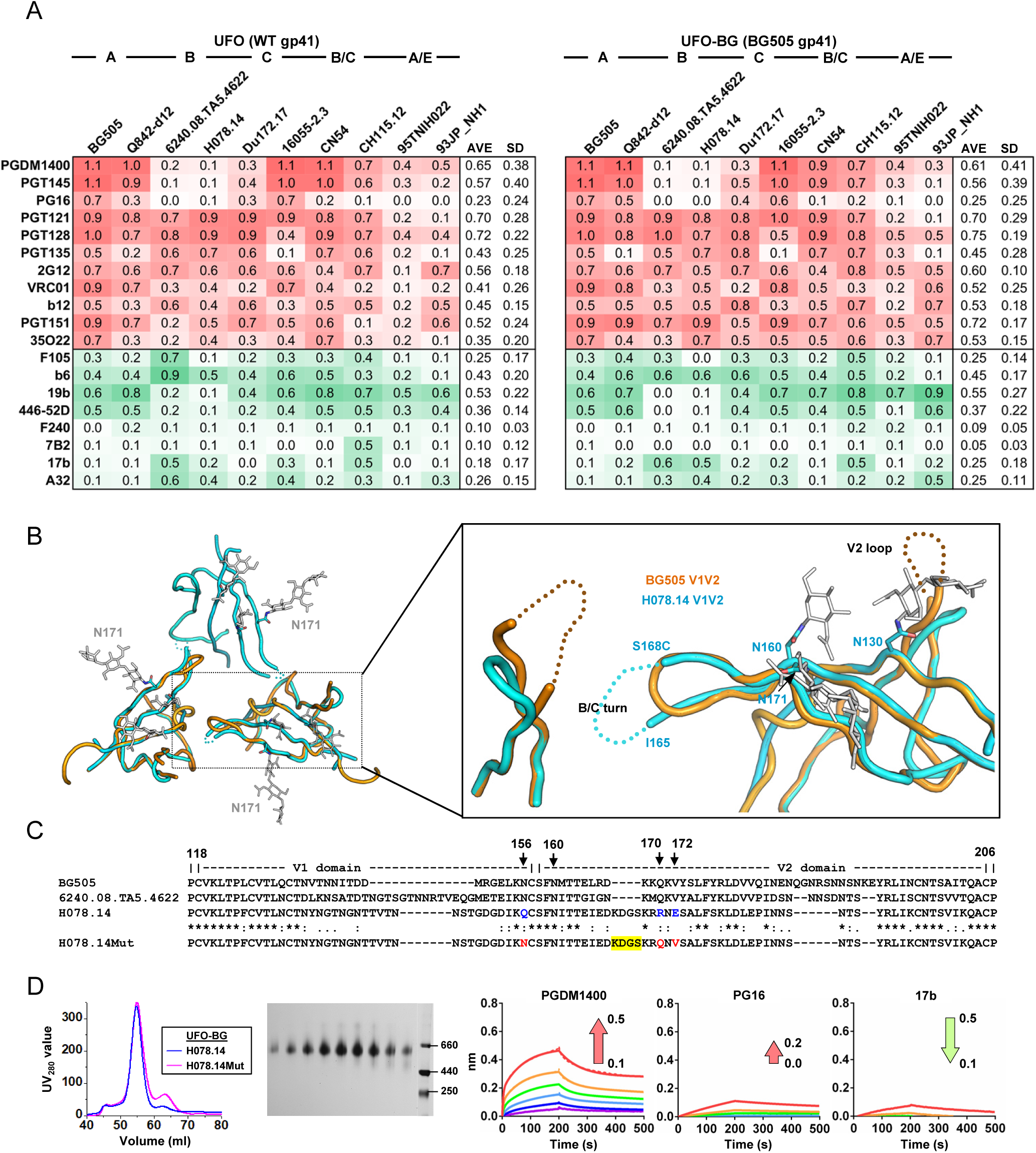
Antigenic map of diverse HIV-1 strains and structure-informed optimization of the H078.14 UFO-BG trimer. (**A**) Antigenic profiles of ten UFO trimers (left) and ten UFO-BG trimers (right) against eleven bNAbs and eight non-NAbs. Sensorgrams were obtained from an Octet RED96 using a trimer titration series of six concentrations (200-6.25 nM by two-fold dilution) and are shown in Figure S4. The peak values at the highest concentration are summarized in the matrix, in which cells are colored in red and green for bNAbs and non-NAbs, respectively. Higher color intensity indicates greater binding signal measured by Octet. To facilitate antigenic comparison between UFO and UFO-BG trimers, average peak value (AVE) and standard deviation (SD) for each antibody are listed in the last two columns of each matrix. (**B**) A top-down view of the H078.14 UFO-BG trimer apex and a zoomed-in view of the H078.14 V1V2 apex superposed with that of the BG505 SOSIP.664 trimer (PDB: 5CEZ). Glycans at N130, N160, and N171 are labeled for H078.14. The turn between strands B and C of H078.14 and the V2 loop of BG505 are shown as dotted lines in blue and orange, respectively. (**C**) Sequence alignment of V1V2 regions from BG505, 6240.08.TA5.4622 (clade B), WT H078.14 (clade B), and a modified H078.14 (termed H078.14Mut) with mutations at positions 156, 170, and 172 colored in red and "KDGS" deletion at the turn of strands B and C highlighted in yellow shade. (**D**) Characterization of an H078.14Mut construct that also contains a disulfide bond (I201C-A433C) to prevent CD4-induced conformational changes. Trimers produced in 200 ml of ExpiCHO cells are characterized by SEC (left), BN-PAGE (middle), and antigenic evaluation against the V2 apex-directed bNAbs PGDM1400 and PG16 and a CD4i-specific non-NAb 17b (right). The direction and magnitude of the change of peak binding signal (nm) are labeled on the sensorgrams of the H078.14Mut UFO-BG trimer with an arrow colored in red and green for bNAbs and non-NAbs, respectively.

UFO and UFO-BG trimers derived from ten strains of five subtypes, twenty in total, were assessed against nineteen antibodies in 380 Octet experiments (**Figure S4**). The peak antibody-binding signals, as well as average and standard deviation for each antibody, were summarized in two matrices corresponding to UFO and UFO-BG trimers, providing by far the most complete antigenic profiles for these HIV-1 subtypes (Figure 4A). Overall, the two UFO designs exhibited comparable antigenic properties with clade-specific patterns. Notably, trimers derived from clade B 6240.08.TA5.4622 and H078.14 were poorly recognized by apex-directed bNAbs, while shielding the immunodominant V3 and gp41 epitopes more effectively than trimers of other clades. However, this reduced non-NAb recognition of the distal V3 and gp41 epitopes was accompanied by enhanced non-NAb binding to the CD4bs and the CD4i epitope, suggesting localized antigenic features specific to these two clade B Envs. The trimers derived from A/E-recombinant strains displayed similar antigenic patterns, with relatively weak binding to most of the antibodies tested here. Based on the peak binding signals averaged across clades, substitution of WT gp41_ECTO_ with BG505 gp41_ECTO_ of the UFO design improved PGT151 and 35O22 binding by over 40%, likely due to a restored quaternary epitope at the gp120-gp41 interface. However, this gp41_ECTO_ swapping exerted a more complicated effect on other bNAb epitopes, causing small variations in peak signal and, in some cases, binding kinetics. For example, for clade A tier 2 Q842-d12, the UFO-BG trimer bound to PGDM1400 and PG16 with a faster association rate than the UFO trimer, whereas the B/C-recombinant CN54 trimer showed a decreased on-rate in PGDM1400 and PGT145 binding after gp41_ECTO_ swapping (**Figures S4B** and **S4G**). Nonetheless, large-scale antigenic profiling by BLI confirmed that UFO-BG trimers present a close antigenic mimic of native Env.

### Structure-Informed Optimization of a Tier 3 Clade B UFO-BG Trimer

Even with a medium-resolution structure (Figure 3A), the H078.14 UFO-BG trimer provides a valuable template for vaccine design, as it only bound to three of the eight non-NAbs (Figure 4A). However, it is imperative to first determine the cause of poor bNAb binding to the V2 apex, which appeared to be inconsistent with the native-like prefusion trimer conformation. To this end, we superposed the apices of H078.14 UFO-BG and BG505 SOSIP trimer structures (Garces et al., 2015) for visual inspection, which revealed a short insertion at the tip of the V2 hairpin, an additional *N*-linked glycan at position N171 (HXB2 numbering), and a shortened V2 loop (Figure 4B). Based on the sequence alignment and the crystal structure, we identified two more residues in strand C that may have destabilized the V2 apex. Specifically, the inward-facing Q170 and V172 in BG505 are now replaced with charged bulky residues R170 and E172 in H078.14 (Figure 4C). Based on this information, we sought to optimize the H078.14 UFO-BG trimer by restoring the V2 apex with a triple mutation in strand C (Q156N/R170Q/E172V, Q156N to restore this *N*-glycosylation site) and a deletion at the tip of the V2 hairpin (ΔKDGS), and by blocking the CD4-induced conformational change with an I201C-A433C disulfide bond (Do Kwon et al., 2015; Guenaga et al., 2015). As shown in SEC and BN-PAGE, the modified H078.14 UFO-BG trimer retained the high yield and high purity of the original construct (Figure 4D). In BLI assays, this trimer was well recognized by PGDM1400, but still less effectively by PG16, while showing significantly reduced binding to a CD4i non-NAb, 17b (Figure 4D). Our analysis thus confirmed that H078.14 can evade recognition by known bNAbs to the apex through mutations in strand C and surrounding loops. The additional glycan at N175 appeared to have little effect on apex bNAb binding as it points sideways in the crystal structure (Figure 4B, left). However, we noted that clade B 6240.08.TA5.4622, which also showed poor trimer binding to apex bNAbs, possesses the same amino acids as BG505 at those positions critical to H078.14 (Figure 4C), suggesting that this tier 2 virus must utilize a different mechanism to shield its apex.

### Envelope Metastability Has an Evolutionary Root

The genetic diversity of HIV-1 has been extensively studied (Korber et al., 2000) and is considered a crucial factor for vaccine design (Gaschen et al., 2002; Nickle et al., 2003). Envs obtained from transmitted/founder (T/F) viruses or derived from a sequence database by phylogeny, which both represent ancestral states of HIV-1, have been evaluated as vaccine immunogens (Gao et al., 2005; Liao et al., 2006; Liao et al., 2013b). The T/F viruses have been found to exhibit greater infectivity relative to chronic viruses (Parrish et al., 2013). Considering that BG505 is a T/F virus (Wu et al., 2006) and BG505 gp41_ECTO_ of the UFO design can stabilize diverse Envs, we hypothesized that the superior infectivity of T/F viruses could be a result of greater gp41_ECTO_ stability, and as such, the T/F UFO trimers might exhibit higher yield, purity, and stability (Figure 5A). To explore this hypothesis, we tested UFO and UFO-BG trimers for three T/F strains: clade B B41 (Pugach et al., 2015; Sanders et al., 2015; Sullivan et al., 2017), clade C CH505 (Bonsignori et al., 2016; Gao et al., 2014; Liao et al., 2013a), and clade C 1086 (Guenaga et al., 2017; Liao et al., 2013b). Following 100-ml ExpiCHO expression and GNL purification, the Env protein was characterized by SEC (Figure 5B). For clade B T/F B41, UFO and UFO-BG trimers showed similar purity, with a greater yield observed for UFO-BG. For two clade C T/F Envs, a notably high trimer yield was observed for the UFO-BG design. B41 UFO and CH505 UFO-BG were then assessed by BLI using a small panel of antibodies, both displaying antigenic profiles consistent with native-like trimers (Figure 5C). In a recent study, Sullivan et al. screened 852 mutations to improve the antigenicity and stability of a B41 SOSIP trimer (Sullivan et al., 2017). In another study, Guenaga et al. screened various glycine substitutions in the HR1 bend in addition to other stabilizing mutations to improve the 1086 NFL trimer (Guenaga et al., 2017). Our results suggest that UFO and UFO-BG may provide a simple and effective alternative to the screening-based approaches for engineering native-like T/F trimers.

**Figure 5.**
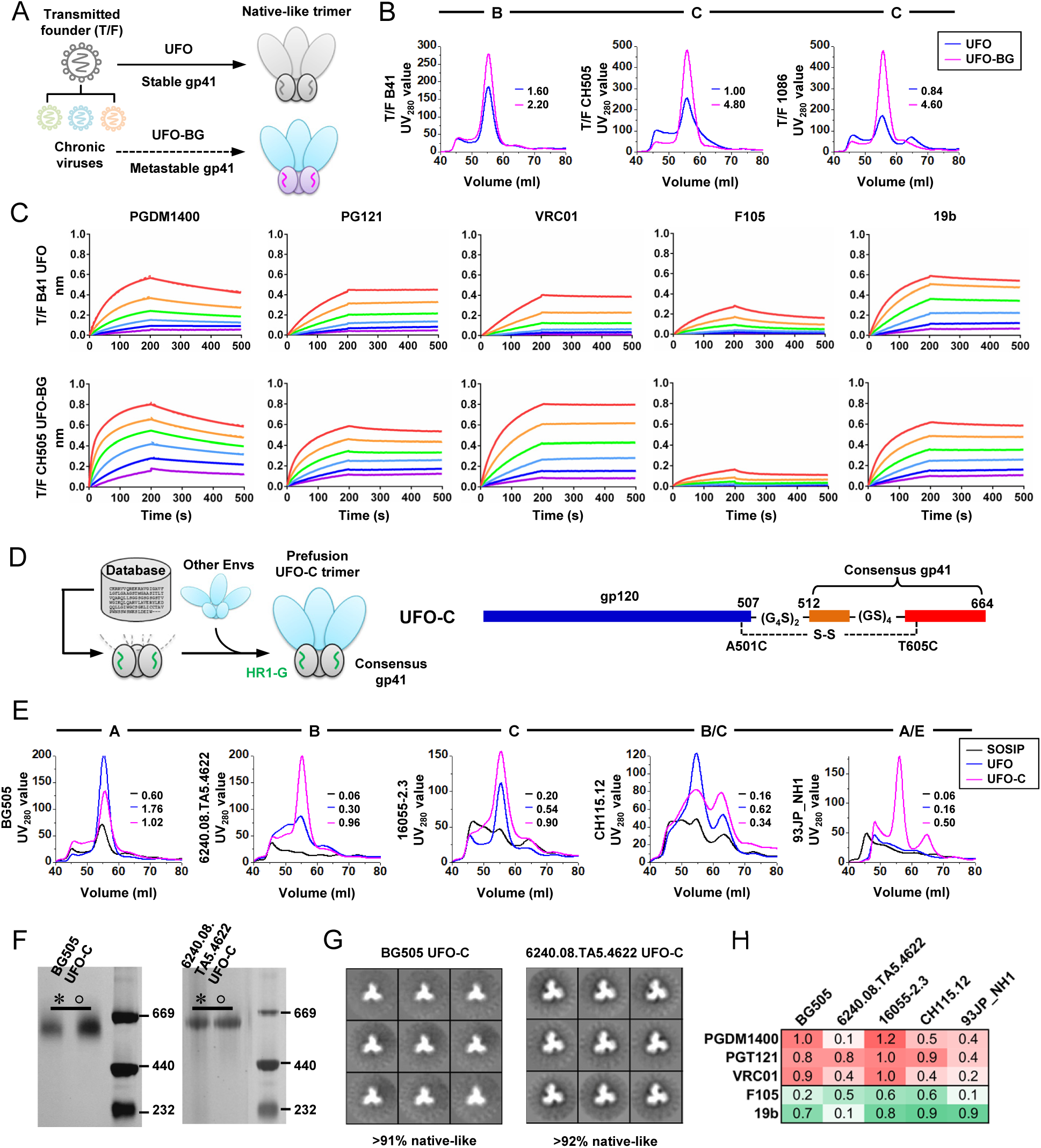
Evolutionary root of Env metastability and design of UFO-C trimers containing a database-derived gp41_ECTO_. (**A**) Connection between HIV-1 evolution and Env metastability. (**B**) SEC profiles of UFO and UFO-BG trimers derived from three transmitted/founder (T/F) viruses, including clade B B41, clade C CH505, and clade C 1086. The yield (mg) of SEC-purified trimer protein (fractions corresponding to 53-57ml) obtained from 100-ml ExpiCHO expression is listed for both trimer designs. (**C**) Antigenic profiles of B41 UFO and CH505 UFO-BG trimers measured against three bNAbs and two non-NAbs. Sensorgrams were obtained on an Octet RED96 using a trimer titration series of six concentrations (200-6.25 nM by two-fold dilution). (**D**) Design (left) and schematic representation (right) of the UFO-C trimers. As shown on the left, a database-derived consensus gp41_ECTO_ in conjunction of a generic UFO design is used to stabilize gp120s from diverse HIV-1 Envs in a hybrid form of gp140 trimer designated UFO-C. The generic GS linker at the HR1 bend is highlighted in green. (**E**) SEC profiles of UFO-C trimers derived from five representative stains of clade A, B, C, B/C, and A/E origins. The yield (mg) of SEC-purified trimer protein (fractions corresponding to 53-57ml) obtained from 100-ml ExpiCHO expression is listed for each of the three trimer designs (SOSIP, UFO, and UFO-C). (**F**) BN-PAGE of Env proteins after GNL purification but prior to SEC and of purified trimers following SEC and BN-PAGE for two UFO-C constructs based on clade A BG505 and clade B 6240.08.TA5.4622. (**G**) Reference-free 2D class averages derived from negative-stain EM of the BG505 and 6240.08.TA5.4622 UFO-C trimers, with the full sets of images shown in Figure S5A. Percentage of native-like trimers is indicated for each trimer construct. (**H**) Antigenic profiles of five purified UFO-C trimers measured against three bNAbs and two non-NAbs. Sensorgrams were obtained on an Octet RED96 using a trimer titration series of six concentrations (200-6.25 nM by two-fold dilution) and are shown in Figure S5B. The peak values at the highest concentration are summarized in the matrix, in which cells are colored in red and green for bNAbs and non-NAbs, respectively. Higher color intensity indicates greater binding signal measured by Octet.

We next explored the UFO design utilizing a consensus gp41_ECTO_ derived from the available Env sequences in the database, designated UFO-C (Figure 5D). If such a UFO-C design is proven successful, it will provide further evidence for the evolutionary root of metastability as consensus has been considered a simple approximation to the ancestral state in HIV-1 evolution (Kothe et al., 2006; Nickle et al., 2003; Santra et al., 2008). To this end, we derived a consensus gp41_ECTO_ from 6,670 full-length Env sequences (https://www.hiv.lanl.gov/) with the HR1 bend replaced by a generic HR1 linker as reported in our previous study (Kong et al., 2016a). The UFO-C constructs were created for five Envs of different subtypes and characterized by SEC following ExpiCHO expression and GNL purification (Figure 5E). Overall, UFO-C outperformed SOSIP and UFO for five and three Envs, respectively, displaying improved trimer yield and purity. For clade A BG505, UFO-C exhibited a SEC profile similar to that of UFO, but with slightly increased aggregates and decreased trimer yield. This result confirmed that consensus gp41_ECTO_ can recapitulate, in large part, the inherent stability of the BG505 gp41_ECTO_. UFO-C appeared to be least effective for a tier 3 B/C-recombinant strain, CH115.12, for which UFO-BG was also less successful than for other Envs (Figure 2B). UFO-C trimers derived from clade A and B Envs were further validated by BN-PAGE and showed no difference in purity before and after SEC (Figure 5F). Consistently, negative-stain EM confirmed that over 90% of UFO-C trimers were native-like (Figure 5G and **Figure S5A**). Lastly, BLI was utilized to assess the antigenicity of five UFO-C trimers (Figure 5H and **Figure S5B**). In general, UFO-C exhibited antigenic profiles on par with UFO-BG for Envs of clades A and B but not others, suggesting that further optimization may be required for consensus gp41_ECTO_ to achieve the same level of stability as BG505 gp41_ECTO_.

### Nanoparticle Presentation of UFO-BG Trimers Derived from Diverse Subtypes

It has been well established that nanoparticle display of antigens elicits stronger immune responses than non-arrayed antigens (Jennings and Bachmann, 2008; Kushnir et al., 2012; Ludwig and Wagner, 2007; Rodriguez-Limas et al., 2013). However, creating trimer-presenting nanoparticles by the gene fusion approach has proven difficult and was only reported for clade A BG505 (He et al., 2016; Sliepen et al., 2015). On the surface of such gp140 nanoparticles, gp41_ECTO_ would form a “neck” region that connects the gp140 trimer and the nanoparticle backbone beneath. Here, we hypothesized that BG505 gp41_ECTO_ of the UFO design can facilitate both gp140 trimerization and nanoparticle assembly (Figure 6A). To validate this hypothesis, we displayed eight UFO-BG trimers of five subtypes on a ferritin (FR) nanoparticle, which was previously used to present an HR1-resesigned BG505 trimer (He et al., 2016). Briefly, UFO-BG-FR constructs were designed by fusing the C terminus of gp41_ECTO_ (residue 664) to the N terminus (Asp5) of a ferritin subunit. These constructs were expressed transiently in 100-ml ExpiCHO cells followed by a single-step purification with a 2G12 affinity column. BN-PAGE displayed a distinctive band of high molecular weight corresponding to well-formed UFO-BG-FR nanoparticles for all eight strains (Figure 6B). Nanoparticle assembly was further confirmed by negative-stain EM, showing a visible core decorated with eight trimer spikes protruding from the nanoparticle surface (Figure 6C and **Figure S6A**). The UFO-BG-FR nanoparticles exhibited greater thermal stability than the respective UFO-BG trimers, with T_m_ ranging from 68°C to 70°C (**Figure S6B**). The antigenicity was assessed for five representative UFO-BG-FR nanoparticles using six bNAbs and four non-NAbs. Overall, particulate display retained, and in some cases enhanced, the native-like trimer antigenicity, showing patterns specific to epitopes as well as HIV-1 subtypes (Figure 6D and **Figure S6C**). For the V2 apex, PGDM1400 bound to all nanoparticles with comparable or notably higher affinity than the corresponding trimers (Figure 4A), suggesting that the displayed trimers have native-like, closed conformations. For clade B tier 3 H078.14, the restored binding to apex bNAbs might be explained by the enhanced stability of V2 hairpin due to the effect of molecular crowding in the presence of neighboring trimers on the nanoparticle surface (Cheung et al., 2005), whereas for Du172.17 and 93JP_NH1 the increased affinity for apex bNAbs was likely a result of avidity. For the N332 supersite and the CD4bs, particulate display exerted a more favorable influence on the H078.14 UFO-BG trimer. For the gp120-gp41 interface, while all UFO-BG-FR nanoparticles retained the trimer binding to PGT151 (Blattner et al., 2014), a cross-clade reduction in 35O22 binding was observed due to the constrained angle of approach (Huang et al., 2014) on the ferritin nanoparticle surface. For non-NAbs, UFO-BG-FR nanoparticles and UFO-BG trimers displayed broadly similar antigenic profiles. Our results suggest that UFO-BG trimers of diverse subtypes can be readily displayed on nanoparticles due to their enhanced gp41_ECTO_ stability, thus providing an effective approach for designing heterologous nanoparticle vaccines.

**Figure 6.**
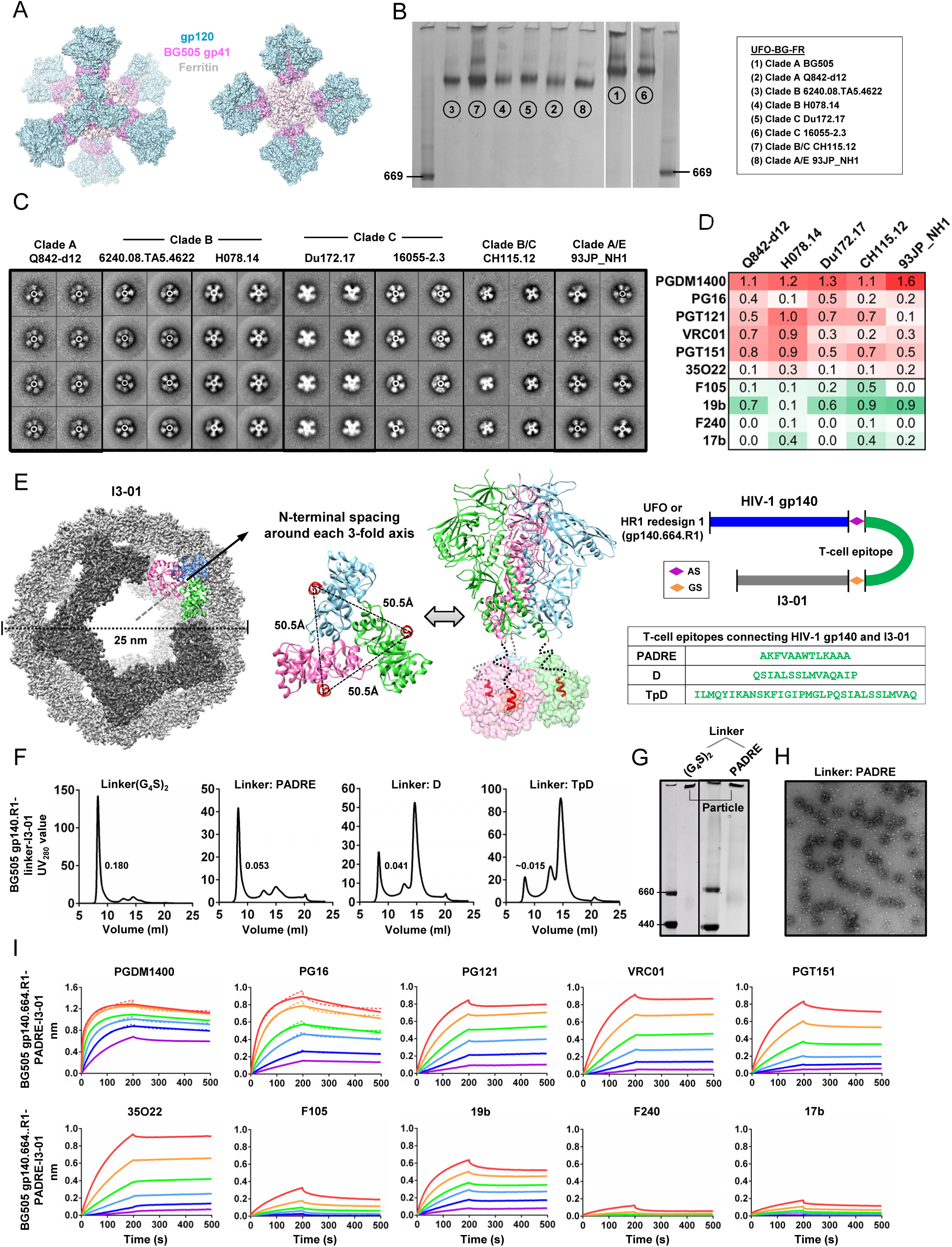
Ferritin nanoparticles presenting diverse UFO-BG trimers and I3-01-based gp140 nanoparticles with embedded T-help signal. (**A**) Surface model of UFO-BG gp140-ferritin (FR) nanoparticle, with gp120, BG505 gp41_ECTO_ of the UFO design, and ferritin colored in cyan, magenta, and gray, respectively. (**B**) BN-PAGE of eight UFO-BG-FR nanoparticles after a singlestep 2G12 affinity purification. (**C**) Reference-free 2D class averages derived from negative-stain EM of five representative UFO-BG-FR nanoparticles. (**D**) Antigenic profiles of five representative UFO-BG-FR nanoparticles against six bNAbs and four non-NAbs. Sensorgrams were obtained on an Octet RED96 using a trimer titration series of six concentrations (starting at 35 nM by two-fold dilution) and are shown in Figure S6. The peak values at the highest concentration are summarized in the matrix, in which cells are colored in red and green for bNAbs and non-NAbs, respectively. Higher color intensity indicates greater binding signal measured by Octet. (**E**) Surface model of the I3-01 nanoparticle (colored in gray) is shown on the left, with the subunits surrounding a frontfacing 5-fold axis highlighted in dark gray and three subunits forming a 3-fold axis colored in sky blue, magenta, and green, respectively. The spacing between N-termini of three I3-01 subunits surrounding a 3-fold axis (top view) and the anchoring of a gp140 trimer onto three I3-01 subunits by flexible linkers (indicated by black dotted lines) are shown in the middle. Schematic representation of I3-01 nanoparticle constructs containing both gp140 and a T-helper epitope is shown on the right, with sequences listed for three such T-helper epitopes, PADRE, D, and TpD. (**F**) SEC profiles of I3-01 nanoparticles presenting an HR1-redesigned BG505 gp140 trimer with a 10-aa GS linker (left) and three different T cell epitope linkers (right). The yield (mg) of 2G12/SEC-purified nanoparticle protein obtained from 100-ml ExpiCHO expression is listed. (**G**) BN-PAGE of two I3-01 nanoparticles, containing a GS linker and a T cell epitope (PADRE) as linker, after a single-step 2G12 affinity purification. (**H**) Micrograph derived from negative-stain EM of an I3-01 nanoparticle presenting an HR 1-redesigned BG505 gp140 trimer with PADRE used as a linker. (**I**) Antigenic profiles of BG505 gp140-PADRE-I3-01 nanoparticle against six bNAbs and four non-NAbs. Sensorgrams were obtained on an Octet RED96 using a trimer titration series of six concentrations (starting at 14 nM by two-fold dilution).

### Design of Trimer-Presenting Nanoparticles with Built-in T Cell Help

Previously, we designed and characterized gp120 and gp140 nanoparticles based on a large 60- mer, E2p (He et al., 2016). Recently, Hsia et al. reported a hyperstable 60-mer (I3-01) resistant to guanidine hydrochloride at high temperature (Hsia et al., 2016). Our database search identified a bacterial enzyme from *Thermotoga maritima* with only five residues differing from I3-01 that has been crystalized at 2.3 Å-resolution (PDB: 1VLW) (**Figure S6D**). Here, we examined the utility of I3-01 for designing gp140 nanoparticles. In terms of symmetry (dodecahedron) and size (25 nm), I3-01 (Figure 6E, left) closely resembles E2p (He et al., 2016). However, the large spacing between the N termini of I3-01 subunits, ~50.5Å, requires a long linker to connect with the C termini of the gp140 trimer (29.1Å) (Figure 6E, middle). We thus hypothesized that a helper T cell epitope may be used not only as a linker between gp140 and I3-01 but also as an embedded signal to boost T cell response and to accelerate B cell development towards bNAbs (Havenar-Daughton et al., 2017). To explore this possibility, we designed three constructs, each containing an HR1-redesigned BG505 gp140 (Kong et al., 2016a), one of the three selected T cell epitopes: PADRE (Alexander et al., 1994), D, and TpD (Fraser et al., 2014), and an I3-01 subunit (Figure 6E, right). A fourth construct containing a 10-aa (G_4_S)_2_ linker was included for comparison. Following furin co-expression in ExpiCHO cells, the 2G12-purified protein was characterized by SEC (Figure 6F). A 10-aa GS linker resulted in I3-01 nanoparticles of high yield and high purity, whereas the three T cell epitopes appeared to affect nanoparticle assembly to various extents due to their hydrophobic nature. Of the three T cell epitopes, PADRE produced nanoparticles of the highest purity, as indicated by SEC (Figure 6F), BN-PAGE (Figure 6G), and negative-stain EM (Figure 6H). In BLI assays, the gp140.664.R1-PADRE-I3-01 nanoparticle exhibited a desirable antigenic profile with strong bNAb binding and minimal non-NAb binding (Figure 6I). To probe the stability of this nanoparticle, we designed ten variants based on the original gene of I3-01, 1VLW (**Figure S6E**). The SEC profiles revealed the importance of a hydrophobic patch at the dimeric interface that facilitates nanoparticle assembly (**Figure S6F**). Taken together, we have re-engineered a hyperstable nanoparticle platform, which displays twenty gp41_ECTO_-stablized trimers on the surface with a built-in T cell help signal.

### Nanoparticles Potently Activate B Cells Expressing bNAbs

Previously, we demonstrated that various BG505 gp120 and gp140 nanoparticles could engage B cells expressing cognate VRC01 receptors (He et al., 2016). Herein, we assessed the degree of B cell activation by five UFO-BG-FR nanoparticles and a BG505 gp140-PADRE-I3-01 nanoparticle with respect to trimers (Figure 7 and **Figure S7A**). B cells expressing bNAbs PGT145, VRC01, and PGT121 (Ota et al., 2012) were used in this assay. Overall, trimer-presenting nanoparticles stimulated bNAb-expressing B cells more effectively than trimers, with peak signals approaching the maximal activation by ionomycin. However, the results also revealed an epitope-dependent pattern: when tested in B cells expressing bNAb PGT121, which recognize the N332 supersite, some trimers and all nanoparticles rendered detectable Ca^2+^ flux signals; in contrast, none and few trimers activated B cells expressing PGT145 and VRC01, which target the V2 apex and the CD4bs, respectively. The stimulation of PGT145-expressing B cells by H078.14 UFO-BG-FR provides further evidence that the apex can be stabilized by neighboring trimers on the nanoparticle surface, consistent with the BLI analysis (Figure 6D). A similar effect was also observed for clade A/E 93JP_NH1 UFO-BG-FR, which bound to PGT121 only weakly by BLI but nevertheless induced a strong Ca^2+^ flux signal in PGT121-expressing B cells, suggesting that cross-linking of B cell receptors (BCRs) by nanoparticles may help overcome the inherent low affinity of trimers. As a result, these nanoparticles will likely elicit more effective NAb response than soluble trimers, thus providing more promising vaccine immunogens.

**Figure 7.**
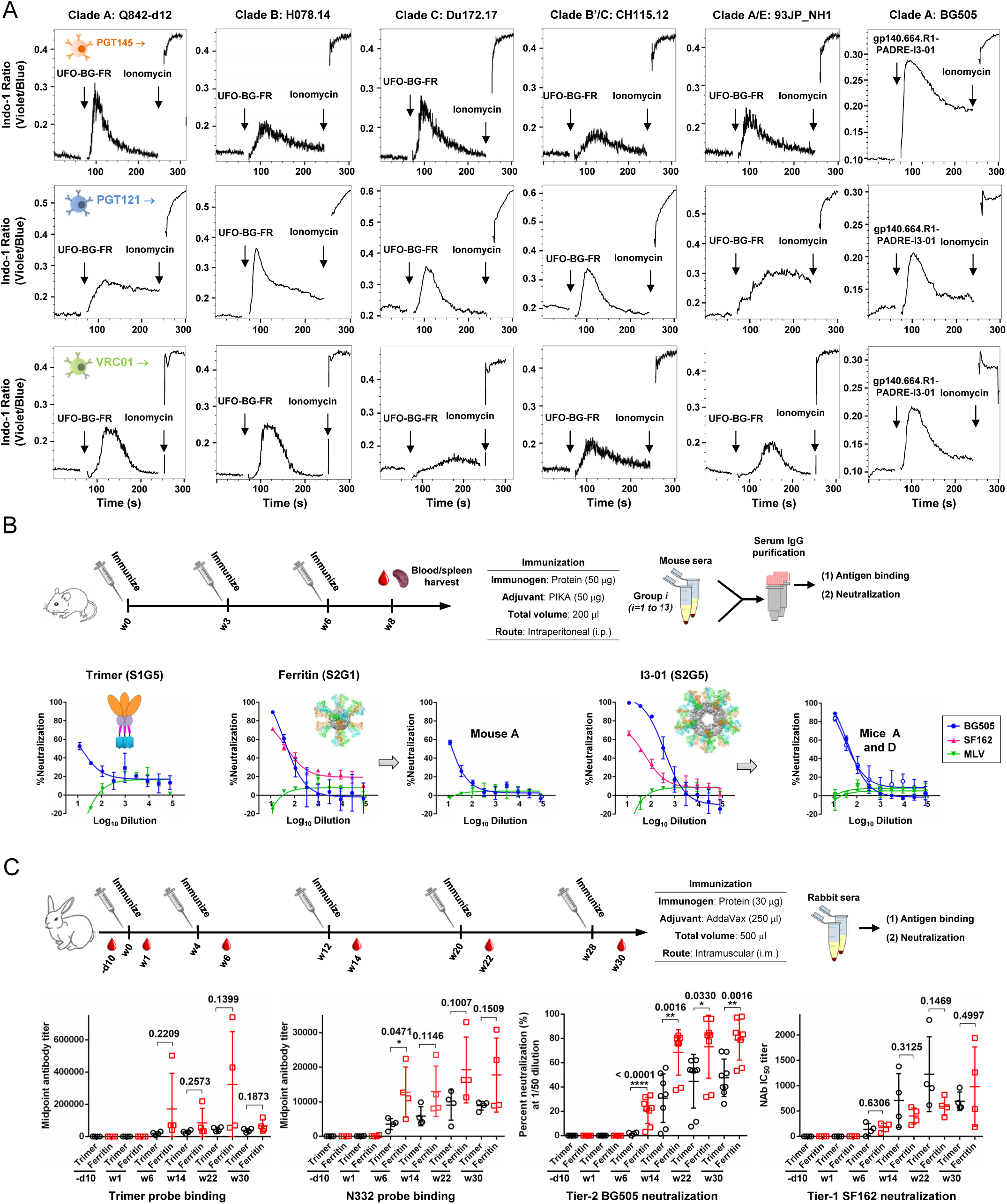
Evaluation of trimers and nanoparticles in B cell activation assays and in two small animal models. (**A**) Ca^2+^ mobilization by various gp140 nanoparticles in B cell transfectants carrying PGT145, PGT121, and VRC01 bNAb receptors. WEHI231 cells expressing a doxycyclin-inducible form of bNAb B cell receptor (BCR) were stimulated with anti-BCR antibodies or the indicated antigens at a concentration of 10 μg ml^-1^: anti-human Ig κ-chain F(ab’)_2_; anti-mouse IgM; an UFO-BG-FR nanoparticle derived from a clade A, B, C, B/C, or A/E strain, or a BG505 gp140-PADRE-I3-01 nanoparticle containing a redesigned HR1 bend within gp41_ECTO_. (**B**) Assessment of immunogenicity in WT mice. Schematic representation of the mouse immunization protocol is shown on the top, and the neutralization curves are shown at the bottom for groups S1G5, S2G1, and S2G5, which correspond to a scaffolded full-length gp140 trimer (gp140.681.R1-1NOG), a gp140-ferritin nanoparticle (gp140.664.R1-FR), and a gp140-I3-01 nanoparticle with T cell help (gp140.664.R1-PADRE-I3-01), respectively. Structural models of three immunogens are placed next to their group-combined neutralization curves. The neutralization curves are also included for individual mice whose serum IgGs neutralized HIV-1. (**C**) Assessment of immunogenicity in rabbits with statistical analysis. Schematic representation of the rabbit immunization protocol (top), longitudinal analysis of midpoint titers of antibodies reactive with the HR 1-redesigned trimer and an N332 nanoparticle probe (bottom, left), and longitudinal analysis of neutralization against autologous tier 2 BG505 and clade B tier 1 SF162 (bottom, right). Percent neutralization (%) at the 50-fold serum dilution and IC50 are plotted for BG505 and SF162, respectively. An unpaired T test was performed to determine whether trimer and ferritin groups were significantly different (P < 0.05) in serum binding and neutralization. P-values are shown for time points w6, w14, w22, and w30, with (^*^) indicating the level of statistical significance. Detailed serum ELISA and neutralization curves are shown in Figures S7D and S7E.

### Rapid Induction of Tier 2 Neutralizing Antibodies in Mice and Rabbits

To obtain an initial readout of immunogenicity, we assessed a subset of gp41_ECTO_-stablized BG505 trimers and nanoparticles in WT mice, where all contained a redesigned HR1 bend – the core of the UFO trimer platform (Kong et al., 2016a). In a previous study, BG505 SOSIP trimers failed to elicit autologous tier 2 NAb response to BG505.T332N in WT mice after eighteen weeks (Hu et al., 2015). It was concluded that the glycan shield of native-like trimers is impenetrable to murine antibodies due to their short heavy-chain complementarity-determining region 3 (HCDR3) loops. Here, we immunized WT BALB/c mice three times with three-week intervals to test eight trimers and four nanoparticles (Figure 7B and **Figure S7B**). PIKA, a toll-like receptor 3 (TLR3) agonist with enhanced T cell and antibody responses reported for a phase-I rabies vaccine trial (Wijaya et al., 2017), was used as an adjuvant. IgG was purified from immunized mouse serum to eliminate non-specific antiviral activity present in the mouse sera (Hu et al., 2015). We assessed the group-combined IgGs with trimer and epitope probes by ELISA (**Figure S7B**). Mouse IgGs elicited by trimers produced in 293 F and ExpiCHO cells (S1G3 and S1G4) bound differentially to the 293 F-expressed probes, supporting our findings in glycan and B cell repertoire analyses (Figures 1D and 1E). The three scaffolded gp140.681 trimers (S1G5, S1G6, and S1G7) elicited stronger antibody responses than the parent gp140.664 trimer (S1G4), consistent with our previous study (Morris et al., 2017). While the ferritin nanoparticle (S2G1) showed a high level of N332-specific IgG response, all three I3-01 nanoparticles (S2G5, S2G6, and S2G7) outperformed the respective trimers containing the same T cell epitopes (S1G8, S1G9, and S1G10). We then evaluated the neutralization of group-combined IgGs (3-8 mg/ml) in TZM-bl assays (**Figure S7C**). Autologous tier 2 neutralization was observed for a scaffolded gp140.681 trimer (S1G5), a ferritin nanoparticle (S2G1), and two I3-01 nanoparticles (S2G5 and S2G6), but not for soluble trimers. When the starting IgG concentration was reduced to 1 mg/ml in neutralization assays, group S1G5 showed a borderline response, whereas one mouse in group S2G1 and two mice in group S2G5 elicited autologous tier 2 NAbs against BG505.T332N at week 8 (Figure 7B). Although IC50 titers cannot be accurately determined for purified IgGs, the results nonetheless suggested a correlation between multivalency (the number of trimers on an antigen) and elicitation of tier 2 NAbs.

Subsequent analysis was then performed on an HR1-redesigned trimer (Kong et al., 2016a) and a ferritin nanoparticle (He et al., 2016) in rabbits with 5 immunizations over 28 weeks (Figure 7C). Rabbit sera collected at six timepoints during immunization were analyzed by ELISA with a trimer probe and an N332 probe (Figure 7C and **Figure S7D**). Overall, rabbits in the trimer group yielded comparable EC_50_ titers that increased steadily over time, whereas rabbits in the ferritin group exhibited a stronger and more rapid antibody response with varying EC50 titers. In the ferritin group, rabbit 65 showed a midpoint titer of trimer-specific response that was 7 to 12-fold greater than others at week 6, and rabbit 63 reached a high level of antibody response at week 22. A similar trend was observed in ELISA binding of the N332 probe: the ferritin group showed mean midpoint 3.6- and 2.1-fold greater than the trimer group at weeks 6 and 22, respectively, with a statistically significant P-value (0.0471) found for week 6. The elevated binding antibody titers might be linked to greater neutralizing activity. To examine this possibility, we conducted TZM-bl neutralization assays longitudinally (Figure 7C and **Figure S7E**). Indeed, the ferritin nanoparticle induced autologous tier 2 NAbs to BG505.T332N more effectively than the trimer, showing a greater percentage of neutralization at the first point of the serum dilution series (50 fold). At week 6, all rabbits in the ferritin group exhibited consistent autologous tier 2 neutralization, whereas the trimer group did not induce such detectable tier 2 NAbs until week 14 (**Figure S7E**). A statistical analysis found that the tier 2 NAb response was significantly different between the ferritin group and the trimer group for week 6 (P<0.0001), week 14 (P=0.0016), week 22 (P=0.0330), and week 30 (P=0.0016). Both rabbit groups showed tier 1 neutralizing activity against clade B SF162 starting at week 6, with greater IC_50_ titers observed for the trimer group, which was consistent with the mouse immunization data (Figure 7B), although this finding is not statistically significant.

Our results demonstrate the superior immunogenicity of nanoparticles displaying gp41_ECTO_-stablized trimers in small animals when compared to trimers alone, and confirm the difficulty of eliciting tier 2 NAbs with soluble trimers independent of the design platform (de Taeye et al., 2015; Klasse et al., 2016; Martinez-Murillo et al., 2017; Sanders et al., 2015). The immunogenicity of gp140 nanoparticles can be improved by optimizing the vaccine regimen (Pauthner et al., 2017), or using large 60-mers designed here and in our previous study (He et al., 2016).

## DISCUSSION

Antibody-based HIV-1 vaccines aim to elicit antibodies capable of neutralizing tier 2 strains, thus preventing the acquisition of infection (Haynes and Mascola, 2017). The advent of the BG505 SOSIP.644 gp140 trimer and structural analysis of trimers based on SOSIP and other designs have established native-like trimers as a promising vaccine platform (Sanders and Moore, 2017; Ward and Wilson, 2017). Accordingly, native-like trimers have elicited autologous tier 2 NAb response in rabbits and NHPs following 6-12 months of immunization (Martinez-Murillo et al., 2017; Pauthner et al., 2017; Sanders et al., 2015; Sliepen et al., 2015). The failure for SOSIP trimers to elicit tier 2 NAbs in WT mice has been attributed to the dense glycan shield and short HCDR3 loops of murine antibodies (Hu et al., 2015). Design of virus-like particles (VLP) presenting native-like Env spikes (Schiller and Chackerian, 2014) and induction of high-quality T follicular helper (Tfh) cells (Havenar-Daughton et al., 2017) were also deemed critical for developing effective HIV-1 vaccine candidates.

Here, we addressed these critical issues in HIV-1 vaccine development with a coherent strategy focusing on Env metastability. Previously, we identified the HR1 bend in gp41_ECTO_ as the primary cause of metastability and developed the UFO platform – containing a redesigned HR1 bend – for trimer stabilization (Kong et al., 2016a). In this study, we demonstrated that gp41_ECTO_ is the main source of metastability by replacing gp41_ECTO_ of diverse Envs with BG505 gp41_ECTO_ of the UFO design and revealed the evolutionary root of metastability by analyzing UFO trimers of three T/F strains and with a consensus gp41_ECTO_. The importance of gp41 stability for nanoparticle vaccines was illustrated by designing ferritin nanoparticles for diverse subtypes and incorporating T cell help into a large 60-mer, which was only possible when gp41_ECTO_ has minimized metastability. A subset of gp41_ECTO_-stablized trimers and gp140 nanoparticles was assessed in mice and rabbits, revealing a critical factor in tier 2 NAb elicitation that has not been apparent or clear in current trimer vaccines: nanoparticles can induce autologous tier 2 NAbs more effectively than trimers with statistical significance (P<0.05), indicating a correlation between antigen valency and tier 2 NAb elicitation. High-quality UFO trimers and nanoparticles can be produced in ExpiCHO cells with substantial yield and purity, bearing important implications for vaccine manufacture.

Future investigation should be directed toward optimization and evaluation of the large, 60-meric I3-01 and E2p nanoparticles, which would be expected to be more effective than the ferritin nanoparticle in the elicitation of tier 2 NAbs. Nanoparticles presenting gp41_ECTO_-stablized trimers of diverse subtypes, when used in a cocktail or sequentially, may be more capable of conferring broad protection against HIV-1.

## EXPERIMENTAL PROCEDURES

### Antibodies

We utilized a panel of bNAbs and non-NAbs to characterize the antigenicity of various native-like trimers and gp140 nanoparticles. Antibodies were requested from the NIH AIDS Reagent Program (https://www.aidsreagent.org/) except for bNAbs PGDM1400, PGT145, PGT121 and PGT151, and non-NAb 19b, which were provided by D.S. and D.R.B.

### Expression and Purification of HIV-1 Env Trimers and Nanoparticles

Trimers were transiently expressed in HEK293 F or ExpiCHO cells (Thermo Fisher) except for crystallographic analysis in which HEK293 F cells were treated with kifunensine. The protocol used for trimer production in HEK293 F cells has been described previously (Kong et al., 2016a; Morris et al., 2017). For cleaved HR1-redesigned trimers, the furin plasmid was added during transfection. The protocol for trimer and nanoparticle production in ExpiCHO cells is as follows. Briefly, ExpiCHO cells were thawed and incubated with ExpiCHO^™^ Expression Medium (Thermo Fisher) in a shaker incubator at 37 °C, with 135 rpm and 8% CO_2_. When the cells reached a density of 10×10^6^ ml^-1^, ExpiCHO^™^ Expression Medium was added to reduce cell density to 6×10^6^ ml^-1^ for transfection. The ExpiFectamine^™^ CHO/plasmid DNA complexes were prepared for 100-ml transfection in ExpiCHO cells following the manufacturer’s instructions. For SOSIP and HR1-redesigned trimers as well as the I3-01 nanoparticles presenting an HR1-redesigned trimer of the BG505 strain, 80 μg of antigen plasmid, 30 μg of furin plasmid, and 320 μl of ExpiFectamine^™^ CHO reagent were mixed in 7.7 ml of cold OptiPRO^™^ medium (Thermo Fisher), whereas for UFO trimers (including UFO-BG and UFO-C) as well as UFO-BG-FR nanoparticles, 100 μg of antigen plasmid was used without furin. After the first feed on day 1, ExpiCHO cells were cultured in a shaker incubator at 32 °C, with 120 rpm and 8% CO_2_ following the Max Titer protocol with an additional feed on day 5 (Thermo Fisher). Culture supernatants were harvested 13 to14 days after transfection, clarified by centrifugation at 4000 rpm for 20 min, and filtered using a 0.45 μm filter (Thermo Fisher). For trimers, Env protein was extracted from the culture supernatants using a *Galanthus nivalis* lectin (GNL) column (Vector Labs), whereas for nanoparticles, Env-fusion protein was purified using a 2G12 affinity column. Some trimers were further purified by size exclusion chromatography (SEC) on a Superdex 200 Increase 10/300 GL column or a HiLoad 16/600 Superdex 200 PG column (GE Healthcare). The purity of I3-01 nanoparticles was characterized by SEC on a Superose 6 10/300 GL column. For both trimers and nanoparticles, protein concentration was determined using UV_280_ absorbance with theoretical extinction coefficients.

### Analysis of Total and Site-Specific Glycosylation Profiles

The total glycan profiles of ExpiCHO and 293 F-produced trimers were generated by HILIC-UPLC. *N*-linked glycans were enzymatically released from envelope glycoproteins via in-gel digestion with Peptide-N-Glycosidase F (PNGase F), subsequently fluorescently labelled with 2-aminobenzoic acid (2-AA) and analyzed by HILIC-UPLC, as previously described (Behrens et al., 2017; Behrens et al., 2016; Neville et al., 2009; Pritchard et al., 2015). Digestion of released glycans with Endo H enabled the quantitation of oligomannose-type glycans (Pritchard et al., 2015). The composition of the glycans was determined by analyzing released glycans from trimers by PNGase F digestion using ion mobility MS (Behrens et al., 2016). Negative ion mass, collision-induced dissociation (CID) and ion mobility spectra were recorded with a Waters Synapt G2Si mass spectrometer (Waters Corp.) fitted with a nano-electrospray ion source. Waters Driftscope (version 2.8) software and MassLynx^™^ (version 4.1) were used for data acquisition and processing. Spectra were interpreted as described previously (Harvey, 2005a, b, c; Harvey et al., 2008). The results obtained served as the basis for the creation of sample-specific glycan libraries, which were used for subsequent site-specific *N*-glycosylation analyses. For site-specific *N*-glycosylation analysis, before digestion, trimers were denatured and alkylated by incubation for 1h at room temperature (RT) in a 50 mM Tris/HCl, pH 8.0 buffer containing 6 M urea and 5 mM dithiothreitol (DTT), followed by addition of 20 mM iodacetamide (IAA) for a further 1h at RT in the dark, and then additional DTT (20 mM) for another 1h, to eliminate any residual IAA. The alkylated trimers were buffer-exchanged into 50 mM Tris/HCl, pH 8.0 using Vivaspin columns and digested separately with trypsin and chymotrypsin (Mass Spectrometry Grade, Promega) at a ratio of 1:30 (w/w). Glycopeptides were selected from the protease-digested samples using the ProteoExtract Glycopeptide Enrichment Kit (Merck Millipore). Enriched glycopeptides were analyzed by LC-ESI MS on an Orbitrap fusion mass spectrometer (Thermo Fisher Scientific), as previously described (Behrens et al., 2016), using higher energy collisional dissociation (HCD) fragmentation. Data analysis and glycopeptide identification were performed using Byonic^™^ (Version 2.7) and Byologic^™^ software (Version 2.3; Protein Metrics Inc.), as previously described (Behrens et al., 2016).

### Blue Native Polyacrylamide Gel Electrophoresis (BN-PAGE)

Env proteins and nanoparticles were analyzed by blue native polyacrylamide gel electrophoresis (BN-PAGE) and stained with Coomassie blue. The protein samples were mixed with G250 loading dye and added to a 4-12% Bis-Tris NuPAGE gel (Life Technologies). BN-PAGE gels were run for 2.5 hours at 150 V using the NativePAGE^™^ running buffer (Life Technologies) according to the manufacturer’s instructions.

### Differential Scanning Calorimetry (DSC)

Thermal stability of UFO-BG trimers, UFO-C trimers, and trimer-presenting nanoparticles was measured using a MicroCal VP-Capillary calorimeter (Malvern) in PBS buffer at a scanning rate of 90 °Ch^-1^ from 20 °C to 120 °C. Data were analyzed using the VP-Capillary DSC automated data analysis software.

### Protein Production and Purification for Crystallization

The clade B tier 3 H078.14 UFO-BG trimer was expressed in FreeStyle 293 F cells treated with kifunensine and purified from culture supernatant using a 2G12 affinity column followed by size exclusion chromatography (SEC). Fabs PGT124 and 35O22 were transiently transfected into FreeStyle 293F cells (Invitrogen) and purified using a LC-λ, capture select column, prior to further purification by ion exchange chromatography and SEC on a Superdex 200 16/60 column. The trimer complexes were prepared by mixing H078.14 UFO-BG trimer protein with PGT124 and 35O22 at a molar ratio of 1:3.5 for 30 min at room temperature. To decrease glycan heterogeneity, deglycosylation was conducted on the PGT124- and 35O22-bound H078.14 UFO-BG Env protein produced in 293 F cells with Endoglycosidase H (New England Biolabs) overnight at 4 °C. The trimer complexes were subjected to crystal trials after further purification of the complexes by SEC.

### Protein Crystallization and Data Collection

The SEC-purified H078.14 UFO-BG trimer complexes were concentrated to ~8 mg/ml before being subjected to extensive crystallization trials at both 4 °C and 20 °C using our automated Rigaku CrystalMation^™^ robotic system (Rigaku) at TSRI (Elsliger et al., 2010). Crystals for protein complex containing Fab PGT124 and 35O22 bound to UFO-BG trimer were obtained from 0.1 Tris (pH 7.4), 0.2 M lithium sulfate, 6% (w/v) polyethylene glycol 4000, using the sitting drop (0.2μl) vapor diffusion method, and harvested and cryo-protected with 30% ethylene glycol, followed by immediate flash cooling in liquid nitrogen. The best crystal diffracted to 4.43 Å resolution and diffraction data were collected at the Advanced Photon Source (APS) beamline 23IDB, processed with HKL-2000 (Otwinowski and Minor, 1997), and indexed in space group P6_3_ with 98% completeness with unit cell parameters *a* = *b* = 127.3 Å, *c* = 316.0 Å (Table S1).

### Structure Determination and Refinement

The H078.14 UFO-BG trimer structure bound to PGT124 and 35O22 was solved by molecular replacement (MR) using Phaser (McCoy et al., 2007) with the BG505 SOSIP.664 protomer:35O22 component from a previous structure (PDB: 5CEZ) (Garces et al., 2015), followed by the PGT124 Fab structure (PDB: 4R26) (Garces et al., 2014) as the molecular replacement models. The structures were refined using Phenix (Adams et al., 2010), with Coot used for model building (Emsley et al., 2010) and MolProbity for structure validation (Chen et al., 2010). Due to the limited resolution of the datasets, two *B*-factor groups per residue were used in refinement. The final R_cryst_ and R_free_ values for complex structure are 32.6% and 34.6%. Figures were generated with PyMol and Chimera (Pettersen et al., 2004). In the crystal structure, the residues were numbered according to the Kabat definition (Martin, 1996) for the Fabs and according to the HXBc2 system for gp140.

### Negative-Stain Electron Microscopy (EM)

UFO-BG trimers, UFO-C trimers, and gp140 trimer-presenting nanoparticles were analyzed by negative-stain EM. A 3 μl aliquot containing ~0.01 mg/ml of the trimers or nanoparticles was applied for 15 s onto a carbon-coated 400 Cu mesh grid that had been glow discharged at 20 mA for 30 s, then negatively stained with 2% (w/v) uranyl formate for 30 s. Data were collected using a FEI Tecnai Spirit electron microscope operating at 120 kV, with an electron dose of ~25 e^-^ Å^-2^ and a magnification of 52,000 × that resulted in a pixel size of 2.05 Å at the specimen plane. Images were acquired with a Tietz 4k × 4k TemCam-F416 CMOS camera using a nominal defocus of 1500 nm and the Leginon package (Suloway et al., 2005). UFO-BG trimer particles were selected automatically from the raw micrographs using DoG Picker (Voss et al., 2009), while trimer-presenting nanoparticles were selected manually using the Appion Manual Picker (Lander et al., 2009). Both were put into particle stack using the Appion software package (Lander et al., 2009). Reference-free, two-dimensional (2D) class averages were calculated using particles binned by two via iterative multivariate statistical analysis (MSA)/multireference alignment (MRA) and sorted into classes (Ogura et al., 2003). To analyze the quality of the trimers (native-like and non-native), the reference free 2D class averages were examined by eye as previously described (de Taeye et al., 2015).

### Bio-Layer Interferometry (BLI)

The kinetics of trimer and nanoparticle binding to bNAbs and non-NAbs was measured using an Octet Red96 instrument (fortéBio, Pall Life Sciences). All assays were performed with agitation set to 1000 rpm in fortéBio 1× kinetic buffer. The final volume for all the solutions was 200 μl per well. Assays were performed at 30 °C in solid black 96-well plates (Geiger Bio-One). 5 μg ml^-1^ of antibody in 1× kinetic buffer was loaded onto the surface of anti-human Fc Capture Biosensors (AHC) for 300 s. A 60 s biosensor baseline step was applied prior to the analysis of the association of the antibody on the biosensor to the antigen in solution for 200 s. A two-fold concentration gradient of antigen, starting at 200 nM for trimers and 14-35 nM for nanoparticles depending on the size, was used in a titration series of six. The dissociation of the interaction was followed for 300 s. Correction of baseline drift was performed by subtracting the mean value of shifts recorded for a sensor loaded with antibody but not incubated with antigen and for a sensor without antibody but incubated with antigen. Octet data were processed by fortéBio’s data acquisition software v.8.1. Of note, for apex-directed bNAbs, experimental data were fitted with the binding equations describing a 2:1 interaction to achieve the optimal fitting results.

### B Cell Activation Assay

Generation of K46 B-cell lines expressing PGT121, PGT145 or VRCO1 has been previously described (Ota et al., 2012). In brief, K46 cells expressing a doxycyclin-inducible form of bNAb B cell receptors (BCRs) were maintained in advanced DMEM (Gibco), supplemented with 10% FCS, Pen/Strep antibiotics, and 2 μg/ml Puromycin (Gibco). Cells were treated overnight in 1 μg/ml doxycyclin (Clontech) to induce human BCR expression. After loading with Indo-1 (Molecular Probes) at 1 μM for one hour at 37°C, washed cells were stimulated with the indicated agents at a concentration of 10 μg ml^-1^: anti-mouse IgM (Jackson ImmunoResearch); UFO-BG or an HR1-redesigned gp140 trimer (Kong et al., 2016a) with a T-helper epitope (PADRE) fused to the C-terminus; UFO-BG-FR or I3-01 nanoparticle presenting an HR1-redesigned gp140 trimer. Calcium mobilization was assessed on a LSR II flow cytometer (BD). In each run, the unstimulated B cells were first recorded for 60 s, then with the testing immunogen added, mixed thoroughly, and recorded for 180 s, followed by addition of 1 μl of 1 μg ml^-1^ ionomycin (Sigma) and recording for another 60 s to verify indo loading.

### Immunization and Serum IgG Purification

Seven-week-old BALB/c mice were purchased from The Jackson Laboratory. The mice were housed in ventilated cages in environmentally controlled rooms at TSRI, in compliance with an approved IACUC protocol and AAALAC guidelines. At week 0, each mouse was immunized with 200 μl of antigen/adjuvant mix containing 50 μg of antigen and 100 μl AddaVax adjuvant (Invivogen) or 50 μl PIKA adjuvant (Yisheng Biopharma) per manufacturer’s instruction via the intraperitoneal (i.p.) route. At week 3 and week 6, the animals were boosted with 50 μg of antigen formulated in AddaVax or PIKA adjuvant. At week 8, the animals were terminally bled through the retro orbital membrane using heparinized capillary tubes. Samples were diluted with an equal volume of PBS and then overlayed on 4.5 ml of Ficoll/Histopaque in a 15 ml SepMate tube (StemCell) and spun at 1200 RPM for 10 min at 20 °C to separate plasma and cells. The plasma was heat inactivated at 56 °C for 1 hr, spun at 1200 RPM for 10 min, and sterile filtered. The cells were washed once in PBS and then resuspended in 1 ml of ACK Red Blood Cell lysis buffer (Lonza). After two rounds of washing with PBS, PBMCs were resuspended in 2 ml of Bambanker Freezing Media (Lymphotec Inc.). Spleens were also harvested and grounded against a 40-μm cell strainer (BD Falcon) to release the splenocytes into a cell suspension. The cells were centrifuged, washed in PBS, treated with 10 ml of RBC lysis buffer as per manufacturer’s specifications, and resuspended in Bambanker Freezing Media for cell freezing. One-third of the total serum per mouse, or 600 μl of serum, was purified using a 0.2-ml protein G spin kit (Thermo Scientific) following the manufacturer’s instructions. Purified IgGs from four mice in each group were combined for characterization by ELISA binding and initial neutralization assays, while purified IgGs from individual mice in two groups, S2G1 and S2G5, were used for further characterization of HIV-1 neutralization in TZM-bl assays. Rabbit immunization and blood sampling were carried out under a subcontract at Covance (Denver, PA). Two groups of female New Zealand White rabbits, four rabbits per group, were immunized intramuscularly with 30 μg of trimer or nanoparticle formulated in 250 μl of adjuvant AddaVax (InvivoGen) with a total volume of 500 μL, at weeks 0, 4, 12, 20, and 28. Blood samples, 15 ml each time, were collected at day −10, weeks 1, 6, 14, 22, and 28, as shown in Figure 7C. Plasma was separated from blood and heat inactivated for ELISA binding and neutralization assays.

### Enzyme-Linked Immunosorbent Assay (ELISA)

Each well of a Costar^™^ 96-well assay plate (Corning) was first coated with 50 μl PBS containing 0.2 μg of the appropriate antigens. The plates were incubated overnight at 4 °C, and then washed five times with wash buffer containing PBS and 0.05% (v/v) Tween 20. Each well was then coated with 150 μl of a blocking buffer consisting of PBS, 20 mg ml^-1^ blotting-grade blocker (Bio-Rad), and 5% (v/v) FBS. The plates were incubated with the blocking buffer for 1 hour at room temperature, and then washed five times with wash buffer. In the mouse sample analysis, purified IgGs were diluted in the blocking buffer to a maximum concentration of 100 μg ml^-1^, whereas in rabbit sample analysis, heat inactivated plasma was diluted by 50-fold in the blocking buffer; both samples were subjected to a 10-fold dilution series. For each sample dilution, a total of 50 μl volume was added to the wells. Each plate was incubated for 1h at room temperature, and then washed 5 times with wash buffer. A 1:2000 dilution of horseradish peroxidase (HRP)-labeled goat anti-mouse or anti-rabbit IgG antibody (Jackson ImmunoResearch Laboratories) was then made in the wash buffer, with 50 μl of this diluted secondary antibody added to each well. The plates were incubated with the secondary antibody for 1 hr at room temperature, and then washed 5 times with wash buffer. Finally, the wells were developed with 50 μl of TMB (Life Sciences) for 3-5 min before stopping the reaction with 50 μl of 2 N sulfuric acid. The resulting plate readouts were measured at a wavelength of 450 nm.

### Pseudovirus Production and Neutralization Assays

Pseudoviruses were generated by transfection of 293 T cells with an HIV-1 Env expressing plasmid and an Env-deficient genomic backbone plasmid (pSG3ΔEnv), as described previously (Sarzotti-Kelsoe et al., 2014). Pseudoviruses were harvested 72 hours post-transfection for use in neutralization assays. Neutralizing activity of purified mouse serum IgGs or heat inactivated rabbit plasma was assessed using a single round of replication pseudovirus assay and TZM-bl target cells, as described previously (Sarzotti-Kelsoe et al., 2014). Briefly, pseudovirus was incubated with serial dilutions of mouse serum IgG or rabbit plasma in a 96-well flat bottom plate for 1 hour at 37 °C before TZM-bl cells were seeded in the plate. Luciferase reporter gene expression was quantified 48-72 hours after infection upon lysis and addition of Bright-Glo^™^ Luciferase substrate (Promega). Data were retrieved from a BioTek microplate reader with Gen 5 software, the average background luminescence from a series of uninfected wells was subtracted from each experimental well, and neutralization curves were generated using GraphPad Prism 6.0, in which values from experimental wells were compared against a well containing virus only. To determine IC_50_ values, dose-response curves were fit by nonlinear regression in Prism. Unpaired T test (non-parametric) in Prism was used to determine whether the two rabbit groups immunized with trimer and ferritin nanoparticle were significantly different (P<0.05) in serum binding and neutralization.

### Mouse Repertoire Sequencing and Bioinformatics Analysis

A 5’-RACE protocol has been developed for unbiased sequencing of mouse B-cell repertoires, as previously described (Morris et al., 2017). Briefly, RNA (including mRNA) was extracted from total PBMCs of each mouse into 30 μl of water with RNeasy Mini Kit (Qiagen). 5’-RACE was performed with SMARTer RACE cDNA Amplification Kit (ClonTech). The immunoglobulin PCRs were set up with Platinum *Taq* High-Fidelity DNA Polymerase (Life Technologies) in a total volume of 50 μl, with 5 μl of cDNA as template, 1 μl of 5’-RACE primer, and 1 μl of 10 μM reverse primer. The 5’-RACE primer contained a PGM/S5 P1 adaptor, while the reverse primer contained a PGM/S5 A adaptor. We adapted the mouse 3’-C _γ_1-3 and 3’-Cμ inner primers as reverse primers for 5’-RACE PCR processing of the heavy chains. A total of 25 cycles of PCR was performed and the expected PCR products (500-600 bp) were gel purified (Qiagen). NGS was performed on the Ion S5 system. Briefly, heavy chain libraries from the same group were quantitated using Qubit^®^ 2.0 Fluorometer with Qubit^®^ dsDNA HS Assay Kit, and then mixed using a ratio of 1:1:1:1 for sequencing. Template preparation and (Ion 520) chip loading were performed on Ion Chef using the Ion 520/530 Ext Kit, followed by sequencing on the Ion S5 system with default settings. The mouse antibodyomics pipeline was used to process the raw data and to determine the distributions of heavy-chain germline gene usage.

## Author Contributions

Project design by L.H., S.K., I.A.W. and J.Z; trimer and nanoparticle construct design by L.H. and J.Z.; protein production, purification, and biochemical characterization by L.H., X.L., and C.J.M.; Env complex crystallization, structure determination, and refinement by S.K., A.S., and I.A.W.; negative-stain EM by J.C., G.O., A.B.W., L.H., and J.Z.; BLI of trimers and nanoparticles by L.H.; mouse serum IgG purification by L.H.; serum-antigen ELISA by L.H., C.J.M., and X.L.; mouse repertoire sequencing by L.H.; mouse serum neutralizing assays by D.S., K.L.S-F., and D.R.B.; rabbit serum neutralization assays by L.H., and Z.J.; manuscript written by L.H., S.K., I.A.W. and J.Z. All authors were asked to comment on the manuscript. The TSRI manuscript number is 29573.

## Acknowledgements

We are very grateful to Dr. M. Elsliger for computer support and H. Tien for crystallization screening. X-ray data sets were collected at the GM/CA@APS-23ID-B beamlines, which have been funded in whole or in part with Federal funds from the National Cancer Institute (ACB-12002) and the National Institute of General Medical Sciences (AGM-12006). This research used resources of the Advanced Photon Source (APS), a U.S. Department of Energy (DOE) Office of Science User Facility operated for the DOE Office of Science by Argonne National Laboratory under Contract No. DE-AC02-06CH11357. Electron microscopy data were collected at the Scripps Research Institute EM Facility. This work was supported by the International AIDS Vaccine Initiative Neutralizing Antibody Center and the Bill & Melinda Gates Foundation through the Collaboration for AIDS Vaccine Discovery (OPP1084519) (A.B.W., I.A.W.), by the Center for HIV/AIDS Vaccine Immunology and Immunogen Discovery (CHAVI-ID UM1 AI100663) (A.B.W., I.A.W.), AI084817 (I.A.W., A.B.W.), AI129698 (J.Z.), and AI125078-01A1 (J.Z.).

## Accession Numbers

Coordinates and structure factors for the complex of clade B tier 3 H078.14 UFO-BG trimer bound to Fabs PGT124 and 35O22 are deposited in the PDB with accession code 6CE0.

